# Brain networks, dimensionality, and global signal averaging in resting-state fMRI: Hierarchical network structure results in low-dimensional spatiotemporal dynamics

**DOI:** 10.1101/229567

**Authors:** Stephen J Gotts, Adrian W. Gilmore, Alex Martin

**Affiliations:** Section on Cognitive Neuropsychology Laboratory of Brain and Cognition, National Institute of Mental Health, National Institutes of Health, Bethesda, MD

**Author notes:** Correspondence to: Stephen J. Gotts, Ph.D., Laboratory of Brain and Cognition, NIMH, NIH, Bldg 10, Rm 4C-217, Bethesda, MD 20892-1366.

## Abstract

One of the most controversial practices in resting-state fMRI functional connectivity studies is whether or not to regress out the global average brain signal (GS) during artifact removal. Some groups have argued that it is absolutely essential to regress out the GS in order to fully remove head motion, respiration, and other global imaging artifacts. Others have argued that removing the GS distorts the resulting correlation matrices, qualitatively alters the results of group comparisons, and impairs relationships to behavior. At the core of this argument is the assessment of dimensionality in terms of the number of brain networks with uncorrelated time series. If the dimensionality is high, then the distortions due to GS removal could be effectively negligible. In the current paper, we examine the dimensionality of resting-state fMRI data using principal component analyses (PCA) and network clustering analyses. In two independent datasets (Set 1: N=62, Set 2: N=32), scree plots of the eigenvalues level off at or prior to 10 principal components, with prominent elbows at 3 and 7 components. While network clustering analyses have previously demonstrated that numerous networks can be distinguished with high thresholding of the voxel-wise correlation matrices, lower thresholding reveals a lower-dimensional hierarchical structure, with the first prominent branch at 2 networks (corresponding to the previously described “task-positive”/“task-negative” distinction) and further stable subdivisions at 4, 7 and 17. Since inter-correlated time series within these larger branches do not cancel to zero when averaged, the hierarchical nature of the correlation structure results in much lower effective dimensionality. Consistent with this, partial correlation analyses revealed that network-specific variance remains present in the GS at each level of the hierarchy, accounting for at least 18-20% of the overall GS variance in each dataset. These results demonstrate that GS regression is expected to remove substantial portions of neurogenic brain signals along with artifacts. We highlight alternative means of controlling for residual global artifacts when not removing the GS.

## INTRODUCTION

Over the past decade, resting-state functional MRI studies have become one of the most utilized approaches to study functional brain organization in humans, as well as to study the physiological bases of common psychiatric and neurological disorders (e.g. Di Martino et al., 2014; Fox & Greicius, 2010; Power et al., 2014b; Raichle, 2015a). However, unlike more traditional task-based fMRI, resting-state studies lack a model of the underlying BOLD signal of interest, complicating the separation of desired signal from a myriad of noise sources known to affect BOLD fMRI measures (see Murphy, Birn, & Bandettini, 2013, for a review). Dozens of studies have now demonstrated the residual presence of relatively whole-brain or “global” artifacts due to influences such as head motion, respiration, and hardware malfunction in the fMRI data even after the application of popular de-noising strategies such as nuisance regression (e.g. Van Dijk et al., 2012; Power et al., 2012; Satterthwaite et al., 2012; see Power et al., 2015, for review) and ICA-based cleaning (e.g. Mowinckel et al., 2012; Satterthwaite et al., 2012). Given the global nature of many of these artifacts and our lack of good simultaneous and independent measures of them, a number of groups have used the global average brain signal (or GS) calculated within a whole-brain mask as a nuisance time series in data cleaning (e.g. Fox et al., 2005; Power et al., 2014a; Satterthwaite et al., 2013). Indeed, removing the GS during nuisance regression does strongly attenuate residual head motion artifacts, as well as respiration-related artifacts, outperforming other nuisance regression strategies to which it’s been compared (e.g. Power et al., 2017b; Power et al., submitted; Satterthwaite et al., 2017).

However, others have argued that the benefit of GS removal comes at a high cost. Murphy et al. (2009) noted that removing the GS re-centers the voxel-wise correlation matrix to 0, resulting in approximately half of the correlation values being negative, even if the original correlations were all positive. Saad et al. (2012; 2013) further pointed out that if GS regression is applied to known (simulated) data, the correlation matrices can be shown to undergo distortion from their true values in a complex manner that depends on the original covariance matrix. This last property has the implication that if comparing two groups of subjects with different covariance/correlation structure (the expected case when conducting a clinical study relative to a control group), the distortions will not be the same in the two groups. Subsequently, several studies have purported to show GS-related distortions of exactly this sort in real datasets (e.g. Gotts et al., 2013b; Hahamy et al., 2014; Yang et al., 2014). Gotts et al. (2013b) made further claims that GS regression can result in poorer experimental validity in the sense that their group comparisons between ASD patients and typically developing controls were no longer in accord with brain-behavior correlations in the ASD group.

Currently, the status of GS regression remains contested and unclear (see Liu et al., 2017, Murphy & Fox, 2017, for reviews). Some groups continue to use it and argue that indeed one must use it in order to satisfactorily remove global artifacts such as head motion (e.g. Power et al., 2014; Satterthwaite et al., 2013). Other groups have discontinued its use, controlling for the contribution of residual head motion and other artifacts through motion matching and group-level covariates in ANCOVA-style analyses (e.g. Gotts et al., 2013b; Saad et al., 2013; Yan et al., 2013). Power et al. (2017a; 2017b) have made the case that while it is true that GS-related distortions can occur in simulated data, these simulations have typically been small-scale and “low dimensional” in terms of the number of distinct networks. If one instead examines simulations with a sufficiently high number of networks that have uncorrelated time series with one another (i.e. “high dimensional”), then the network-specific time series can cancel to near 0 when averaged into a GS, leaving only prominent global artifact time series that are shared across all locations. Under these circumstances, the data distortions due to GS regression can be so small that they are effectively negligible. We concur with Power, Petersen and colleagues that data dimensionality is the critical issue here and will likely determine whether the benefit of GS regression is worth the potential cost.

In the current paper, we set out to characterize the dimensionality of resting-state fMRI time series data using principal component analysis (PCA) in two independent resting-state fMRI datasets (Set 1: N=62, Set 2: N=32). PCA is a multivariate technique that decomposes time series data into a mutually orthogonal (i.e. uncorrelated) set of time series that are ordered by amount of variance explained (Pearson, 1901; Hotelling, 1933; see Jolliffe, 2002, for review), and it is commonly used to assess data dimensionality and for data reduction/compression. We further examine how the PCA view of the data relates to the “network view” of the data using cluster analyses, with network parcellations that are based on thresholding the voxelwise correlation matrices (e.g. Yeo et al., 2011; Power et al., 2011). Finally, we directly examine the question of whether these network-specific time series average away to zero in the GS using partial correlation analyses.

## RESULTS

Before examining actual resting-state fMRI data, we present two simulations for which all of the details are known in order to introduce our main analysis techniques. In *Simulation 1*, we compare 1000 random simulated time series with and without a lower-order correlational structure (either two large clusters of time series or none). We use PCA to examine the variance/covariance of these simulated datasets, further checking the structure using Multidimensional Scaling (MDS) analysis (e.g. Cox & Cox, 2001) - a technique that can compress the similarities of all of the time series into two dimensions for easier viewing. We then examine how the variance/covariance of these datasets are affected when the time series are averaged into a single global time series (the GS). In *Simulation 2*, we examine all of these same features for a dataset with a more complex hierarchical correlational structure (5 large time series clusters, each of which are further divided into 5 smaller clusters for a total of 25 clusters).

### Simulation 1: Two versus zero clusters

In this simulation, we generated 1000 random time series from a Gaussian distribution (mean=0, SD=1), each of length 200 time points (see Figure 1A for example time series and *Methods* for details). In one condition (Two Clusters), we created a simple correlation structure by adding a common random time series to 500 of the original time series and a different common time series to the remaining 500, resulting in two large time series clusters (Figure 1B, left panels). In the other condition (Zero Clusters), no shared time series were added (Figure 1B, right panels). In the case of Two Clusters, a scatterplot of the 1000 time series after MDS analysis nicely shows two discrete groupings of points, green (Cluster 1: time series 1-500) and black (Cluster 2: time series 501-1000). In contrast, the corresponding scatterplot for the Zero Cluster case shows full intermixing of the same time series ranges (green: 1-500, black: 501-1000) in a ring around the center (0,0). When applying PCA to these data, the data are decomposed into a series of mutually orthogonal time series (i.e. with inter-correlations of 0) referred to as “principal components” (PC), with the first PC explaining the most variance, the second PC explaining the next most variance that is uncorrelated with the first, and so on. After applying PCA to the 1000x200 time series data in both cases, Figure 1C shows information about the dimensionality of the time series. The left panel shows the variance of the full time series data that is explained by each individual PC, usually referred to as a “scree plot” of the eigenvalues (e.g. Cattell, 1966; see Jolliffe, 2002, for discussion). Each eigenvalue represents the variance of its corresponding PC. After dividing by the sum of all of the eigenvalues (y-axis of left panel), this gives the proportion of the total variance explained by each individual PC. The right panel, in contrast, shows how much of the full time series data are explained in a cumulative fashion when including the first *X* PCs (e.g. the first 2 PCs in the Two Cluster case explain a combined 50% of the total time series variance).

**Figure 1.**
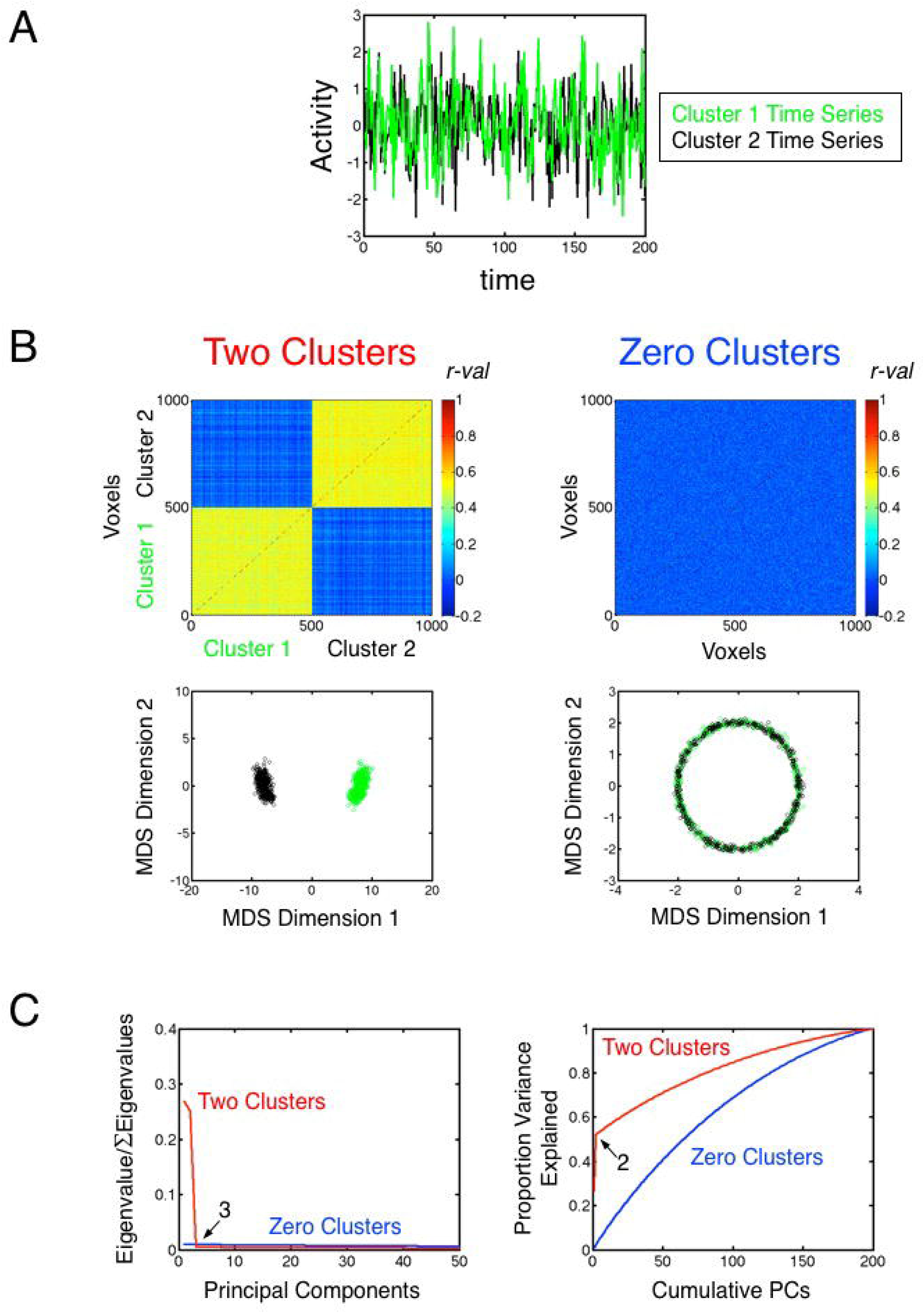
Simulation 1. (A) Sample random time series from the two clusters in the Two Clusters condition. The sample Cluster 1 time series is shown in green, and the sample Cluster 2 time series is shown in black. (B) Top panels show the all-to-all correlation matrices for the 1000 time series, with the Two Clusters condition shown on the left and the Zero Clusters condition shown on the right. Correlation values are indicated by color, with colorbars shown to the right of each plot. The bottom panels show the corresponding two-dimensional MDS scatterplots for the two conditions. The Two Clusters condition on the left shows two well-isolated clusters of points, with each point representing a single time series and Cluster 1 points (time series 1-500) shown in green and Cluster 2 points (time series 501-1000) shown in black. The same ranges of time series for the Zero Clusters condition are shown to the right, with the time series forming a ring around the origin (0,0), as is common in metric MDS for unstructured data (e.g. Hughes & Lowe, 2002). (C) Scree plot of the eigenvalues in the left panel shows proportion of time series variance explained by each individual component in PCA for the two conditions (Two Clusters shown in red, Zero Clusters shown in blue), with the corresponding cumulative time series variance explained by PCs 1-*X* shown in the right panel (where *X* is the component number plotted along the x-axis).

What can one then say about the dimensionality of these cases? In the case of Two Clusters, the scree plot in the left panel shows that the first two PCs are the only ones that explain a large proportion of variance, leveling off to approximately a flat line after this representing a Gaussian “noise tail”, with each added PC only explaining a tiny fraction of the variance. This is expected in this case because there is no easy shortcut to re-representing what are otherwise Gaussian random time series (with correlations near zero before adding the shared time series) than to add approximately one PC for each random time series - as this is the basis of all of the within-cluster variability that remains. Indeed, in the case of Zero Clusters, all of the variance must be explained in this way. Accordingly, the right panel shows a relatively linear increase in variance explained up until the point that 100% of the variance can be explained (i.e. when the number of PCs equals the length of the time series, 200). When using PCA to perform data compression/reduction, a common recommendation is to find the leveling off point or “elbow” (e.g. Thorndike, 1953) of the eigenvalues in the scree plot (3 PCs for Two Clusters) and retain all of the PCs above this point (i.e. PCs 1-2). Other commonly recommended criteria include keeping PCs with raw eigenvalues above 1 (specifically for PCA on correlation matrices; Kaiser, 1960) or requiring that a certain fixed amount of cumulative variance be explained (e.g. 90% of all variance) (see Jolliffe, 2002, for review/discussion). Using the cumulative variance explained criterion can be more prone to inaccuracies when long noise tails are present and the overall signal to noise ratio of the data is low. In the current paper, we will examine both the scree plots of the eigenvalues, as well as the cumulative variance explained, in order to gain a fuller perspective on the dimensionality of the signals involved.

Both of these cases are “high-dimensional” if the goal is to explain most of the unaveraged time series variance in the sets, requiring more than 100 PCs to reach 90% of the variance. However, the Two Clusters case has prominent low-dimensional aspects, with the first two PCs explaining approximately half of all of the variance. If the goal instead is to understand the dimensionality that is most relevant for the GS, then the answer is more unequivocal. When the 1000 time series are averaged into a GS in the Two Clusters case, the Gaussian random variation that determines all of the within-cluster variation averages away to near 0 (since each sample time point was drawn from a distribution with 0 mean and standard deviation of 1), leaving the prototype time series that was added to each cluster (see Figure S1 for a graphical depiction of the Cluster 1 data). Each prototype cluster time series then makes up approximately 50% of the GS, explaining near 100% of the GS variance when combined. In other words, the dimensionality relevant for the GS is 2 rather than 100+, which was nicely conveyed by the scree plot of the eigenvalues. In the case of Zero Clusters, the means of each time point are approximately zero, leaving something close to a flat line in time (see Figure S1). If we were to add even a tiny common signal to all of the time series in the Zero Clusters case, this would come to dominate the GS, since everything else would average away to zero.

### Simulation 2: Hierarchical correlation structure plus a common shared signal

Given the context of *Simulation 1*, we can now consider a slightly more complicated situation. As in the last simulation, each simulated time series (length 200 time points, 1250 total time series) was constructed by first sampling a Gaussian distribution with a mean of 0 and a standard deviation of 1. Five large-scale clusters or “networks” were then defined by adding 5 new random prototype time series (mean=0, SD=1) to different ranges of the 1250 time series (cluster 1: 1-250, cluster 2: 251-500, etc.). Hierarchical structure was then created by further adding five new random time series (mean=0, SD=1) to different ranges of each large-scale network (e.g. for cluster 1, one new random time series was added to series 1-50, another to 51-100, etc.). This resulted in a total of 25 smaller networks that were embedded in the 5 larger networks (see correlation matrix in Figure 2A). Finally, a common random time series was added to all 1250 time series, with a reduced weight of 0.5 (mean=0, SD=0.5). After MDS, a scatterplot of the similarity of the 1250 time series revealed 5 distinct clusters (Figure 2B), corresponding to the five larger “parent” networks, while the distinctions among the 25 finegrained “child” networks were less obvious when viewed in two dimensions. A hierarchical cluster tree was then constructed from the all-to-all correlation matrix using Matlab’s *linkage* Dimensionality in Resting-State fMRI 11 function and the Euclidean distances among the matrix columns (see Figure 2C), which nicely re-capitulated the planned 5x5 network structure. When PCA is applied to the 1250×200 time series dataset, the scree plot of the eigenvalues shows markers of this hierarchical structure (Figure 2D, left panel), with leveling-off points at 6 and 26 PCs. When viewing the cumulative variance explained (Figure 2D, right panel), changes in slope are clear at PCs 5 (46.9% of the variance) and 25 (74.2%), with the noise tail continuing on until 200 PCs and reaching 90% variance explained by 104 PCs.

**Figure 2.**
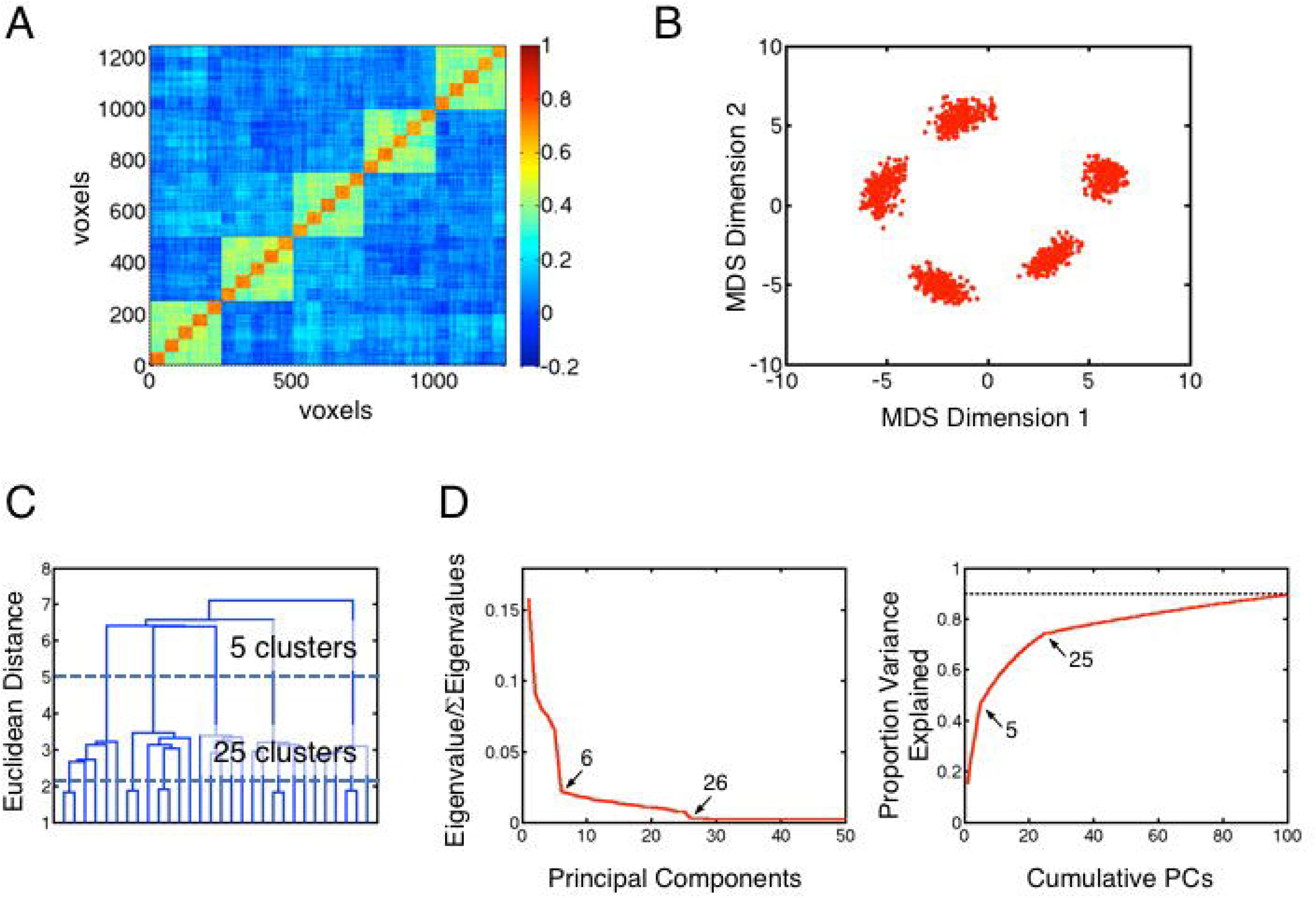
Simulation 2. (A) All-to-all correlation matrix of 1250 random time series with hierarchical network structure added. Perason *r*-values are indicated by color, with colorbar to the right. (B) Two-dimensional MDS scatterplot showing five nicely isolated clusters of points, corresponding to the 5 large scale clusters that make up the first level of the hierarchical structure, with points representing the individual time series out of the 1250. (C) Dendrogram showing the hierarchical cluster tree of the all-to-all correlation matrix, with 5 large clusters that are each broken down further into 5 subnetworks for a total of 25 networks. (D) Scree plot of the eigenvalues showing leveling-off points at PCs 6 and 26 marking aspects of the hierarchical structure (left panel), with corresponding cumulative time series variance explained in the right panel.

As with the previous simulation, the dimensionality relevant for the GS is strongly influenced by the structure of the large-scale networks. When taking only the 5 prototype large network time series (a “coarse” network identification scheme) and entering them into a multiple regression to explain the GS, one can account for 41.2% of the variance in the GS (~ 8% for each large network). Repeating this instead with all 25 subnetwork time series (as one might in a more highly thresholded network identification approach), one reaches 57.0% of the variance in the GS, with 15.8% attributable selectively to the subnetworks beyond the 5 large networks. Further adding the common random signal to the regression model, which might be thought of as a global nuisance signal or a non-specific neurogenic signal, one can account for 99.8% of the variance in the GS (42.8% selectively attributable to the common signal), with only 0.17% of the variance attributable to unaveraged Gaussian variation in the individual time series. If one repeats this simulation, leaving out the 5 large-network time series (resulting in 25 networks, but without a clear hierarchical structure), the variance in the GS selectively explained by the common signal shifts from 42.8% up to 75.2% and the total amount of variance explained by the network structure shrinks from 57% to below 25% (see Figure S2 for a graphical depiction). From these results, it is apparent that one can’t simply look to the number of distinguishable networks in the data (N=25 in both cases) to understand the impact of GS averaging; one must understand and account for the full hierarchical network structure. The presence of lower-order correlation structure can shift the effective dimensionality from high to low because these prominent sources of variance are shared over a wide scale, accounting for large portions of total variance.

For Simulation 2, it is further interesting to note the relationship between the GS and the first PC of the data. Carbonell et al. (2011) have previously argued that extremely high agreement between the GS time series and that of the first PC indicates that no network structure is contaminating the GS. In other words, they argue that the GS represents a linearly additive signal that can be safely removed, with all of the desired network structure contained in PCs 2 and beyond. To examine this issue, we re-ran this simulation 1000 times, each time finding the correlation between the first PC of the 1250×200 time series dataset and the corresponding GS time series. As in the Carbonell et al. (2011) results for the actual fMRI data, the mean correlation between the first PC and the GS across iterations was found to be 0.989 (SD=0.0094), despite the fact that more than 50% of the variance in the GS was due to network-specific sources (mean=55.1%, SD=4.7%). It appears then that Carbonell et al. (2011) may have dismissed network-specific contributions to the GS prematurely.

### PCA analyses of single-subject and group-level fMRI data

Having introduced our analysis tools in the context of *Simulations 1* and *2*, we are now ready to examine the dimensionality of the actual resting-state fMRI data. Since the GS is usually calculated over a whole-brain mask from the single-subject data after the point of volume registration of the individual TRs (i.e. individual EPI time samples of a whole-brain volume) and prior to any nuisance regression (e.g. Fox et al., 2005; Power et al., 2014a), it is the dimensionality of the volume-registered data that is relevant for the GS averaging debate. Two sets of fMRI resting-state data were used for these analyses, one using more traditional single-echo BOLD acquisition (Set 1, N=62) and another using a newer multi-echo acquisition (Set 2, N=32) with a middle echo (TE=27.6 ms) that could be analyzed in the same manner as the single-echo data in Set 1 (TE=27 ms) (see Methods for complete details). Initial preprocessing steps for individual subjects in both datasets involved despiking, correction for slice-time acquisition, volume registration, spatial blurring (6mm FWHM), normalization to percentage signal change, and spatial transformation to Talairach space at a slightly down-sampled spatial resolution (6mm isotropic) to simplify the computational demands of voxelwise analyses for PCA and MDS. Finally, these volume-registered data were baseline detrended (fourth order polynomial model) to remove slow scanner drift in the BOLD time series. The GS was then calculated as the average time series within a whole-brain mask for each subject.

Figure 3A and B (left panels) show the scree plots of the eigenvalues for the individual-subject time series datasets for Set 1 and Set2, respectively. Similar to the examples shown in *Simulations 1* and 2, only the first few eigenvalues explain substantial amounts of variance in terms of individual PCs, with the eigenvalues leveling off prior to 10 PCs in both sets and the most prominent bend in the curves near 4-5 PCs. Correspondingly, the middle panels of Figure 3A and B show a rapid rise in the cumulative variance explained up until around 10 PCs, after which the slopes level off in a more linear fashion until approximately 90 percent of the variance is explained at 50 or more PCs for the typical subject (the mean curve reaches 90% at 56 PCs in Set 1 and 63 PCs in Set 2). Note that these numbers on cumulative variance explained are similar to those observed in some prior studies using PCA to estimate dimensionality of resting-state data (e.g. Kundu et al., 2013, Supporting Information).

**Figure 3.**
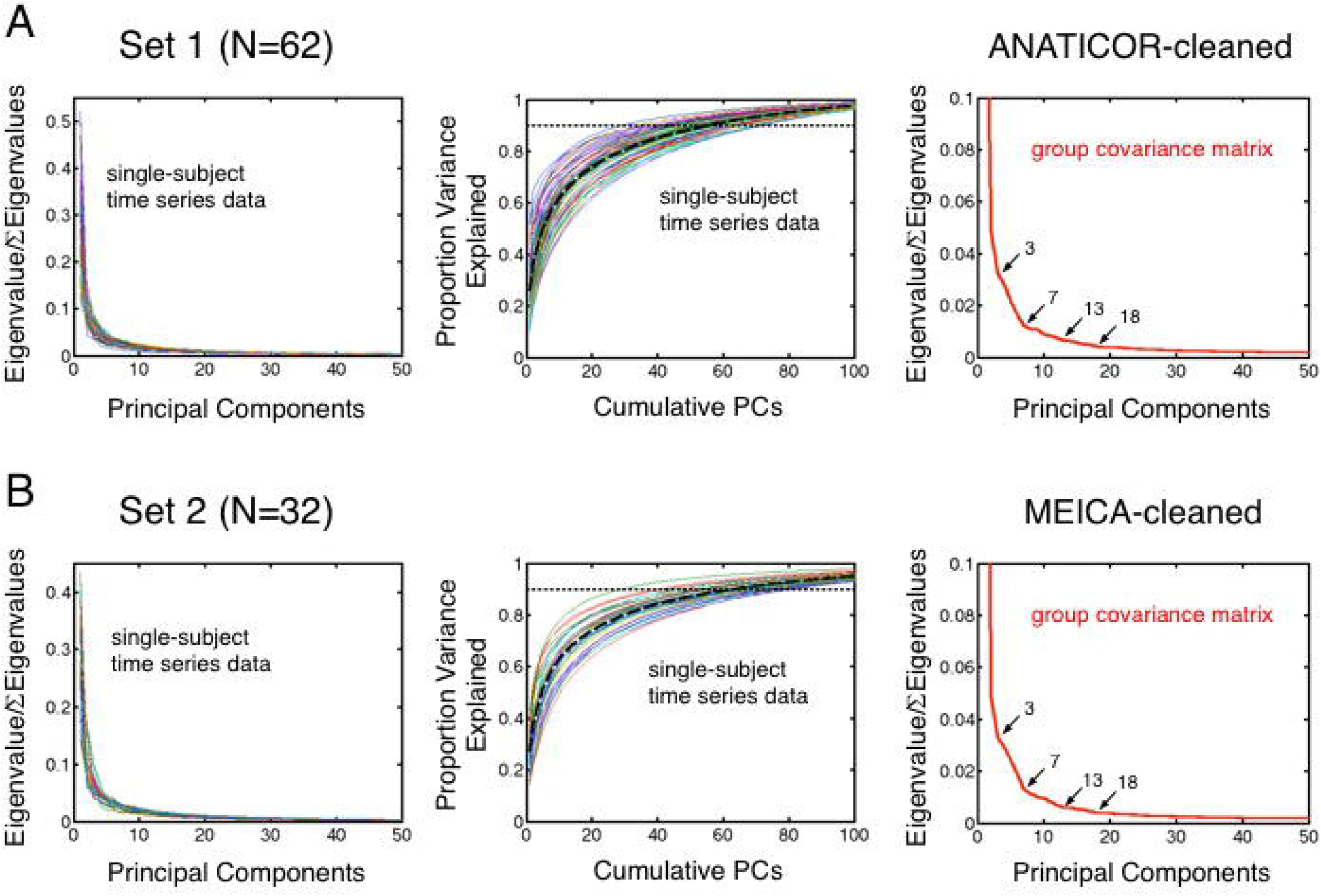
PCA applied to the resting-state time series data for Sets 1 and 2. Scree plots of the eigenvalues for the volume registered data of all individual subjects are shown in the left panels (Set 1 in A, N=62; Set 2 in B, N=32). Middle panels show the corresponding cumulative time series variance explained for the individual subjects, along with the means of these curves in thick dashed black lines. Horizontal dotted lines marking 90% of the variance explained in each set are shown for reference. The right-most panels show similar scree plots of the eigenvalues for the group-average covariance matrices when applying PCA to the de-noised data, using ANATICOR to clean the Set 1 data (A) and MEICA to clean the Set 2 data (B). Prominent leveling-off points that are common to the two sets are marked at 3 and 7, with more minor leveling-off points marked at 13 and 18.

To gain a better understanding of how these PCA results may relate to network structure that is typically studied in de-noised data, we also calculated scree plots on the group average covariance matrices for both sets after de-noising. For Set 1, we used the ANATICOR nuisance regression method that includes modeling physiological artifacts due to cardiac and breathing cycles (Jo et al., 2010, 2013; Gotts et al., 2012), and for Set 2, we utilized a newer method that makes use of the multi-echo features of the data to sort BOLD from non-BOLD variance with ICA (multi-echo ICA or MEIČA; Kundu et al., 2012) (see Methods). We also focused on the gray-matter-only field of view used in prior cortical parcellations with large datasets (Yeo et al., 2011). As shown in the right panels of Figure 3A and B, notable bends in the scree plots of the eigenvalues calculated on the de-noised data are common to both datasets at 3 and 7 PCs, with the sharpest elbow at 7PCs and more minor leveling-off points at 13 and 18 PCs (analogous plots for *Simulations 1* and *2* are shown in Figure 1C and Figure 2D). In the next section, we will examine to what extent these common bends might reflect markers of hierarchical network structure.

### Network clustering analyses reveal low-dimensional hierarchical structure

Previous studies from several labs have already thoroughly characterized the organization of the cerebral cortex into fine-grained network parcellations based on both resting-state and task-based data (e.g., Craddock et al., 2012; Doucet et al., 2011; Gordon et al., 2016; Power et al., 2011; Shen et al., 2013; Shirer et al., 2012; Yeo et al., 2011). Some of these have also previously noted the hierarchical nature of these relationships when thresholding the brain-wide correlation matrices at progressively higher levels (e.g. Power et al., 2011; Yeo et al., 2011; see also Doucet et al., 2011). In these analyses, we conduct parcellations of the cortex in both resting-state datasets using much lower thresholding to examine the lower-dimensional correlation structure. We also examine how this organization relates to previously published finer-grained parcellations that have used large datasets with replication, focusing in particular on the Yeo et al. (2011) 7-network and 17-network parcellations and comparing to views of the voxelwise correlation matrices in the absence of any thresholding using MDS. Finally, we conduct more direct hierarchical clustering analyses using the Yeo et al. (2011) parcellations.

Group-averaged, voxelwise correlation matrices were thresholded at a wide range of levels in both datasets, from as low as the top 90% of the rank-ordered correlation values (referred to as “tie density”) up to the top 0.5% (see Methods for complete details). Two different algorithms were used for cluster analyses: Louvain Modularity (e.g. Blondel et al., 2008; Rubinov & Sporns, 2010) and InfoMap (Rosvall & Bergstrom, 2008, 2011). Louvain Modularity was used for the entire range of thresholds, crossed with several different values of the resolution parameter ‘gamma’, and InfoMap was used for tie densities of 20% and higher. To minimize the inclusion of small and singleton clusters, we required that clusters be at least 1% in size of the total number of voxels being parcellated in order to be retained. Solutions were considered stable when both datasets gave matching cluster solutions at the same or adjacent thresholds, and stable solutions typically spanned a range of adjacent thresholds. At the lower thresholds using the classic modularity metric (gamma=1, 80-90% tie density), stable solutions were observed at N=2 networks, corresponding to the well-known “task-positive”/“task-negative” parcellation (e.g. Fox et al., 2005; shown in yellow and red, respectively, in Figure 4, top two rows), as well as over a wide range of thresholds when using a gamma less than one (0.75). As the thresholds were raised to higher levels, a fronto-parietal network (shown in orange) partially overlapping the first two split off, resulting in N=3 networks, which was seen using both Louvain Modularity (30-50% thresholds) and InfoMap (20% threshold). This was followed by the separation of the remaining task-positive voxels into visual (purple) and somatomotor networks (blue) (N=4 networks, see middle rows of Figure 4), observed for both datasets with Louvain Modularity from 3-20% tie density and with InfoMap for Set 1 from 8-10% tie density. Higher thresholding led to finer-grained parcellations similar to the 7-network and 17-network parcellations of Yeo et al. (2011). However, the solutions were not identical across the two datasets at any higher threshold (<3% tie density) for either Louvain Modularity or InfoMap, likely because our sets involved many fewer subjects than Yeo et al. (2011) and had slightly different scanning fields of view (FOV), which in turn would be expected to lead to less stable estimates of the group-average correlation matrices. For finer-grained parcellations in the current paper, we will therefore use the previously published 7-network and 17-network parcellations of Yeo et al. (2011) that have well-established replication (shown in the bottom two rows of Figure 4). Our datasets do appear to contain these network distinctions, as well, insofar as the correlations within network are greater than the correlations across networks for both parcellations. For Set 1, within-network correlations were greater than the average across-network correlations for all 7 networks [*t*(61)>3.09, *P*<.003 for all 7 comparisons] and all 17 networks [*t*(61)>5.92, Pc.0001 for all 17 comparisons]. For Set 2, within-network correlations were greater than the average across-network correlations for all but the “limbic” network for the 7-network parcellation (shown in white) [*t*(31)>13.92, *P*<.0001 for 6/7 tests; *P*>.4 for limbic network test], which was partly excluded from the field of view for these scans (ventromedial prefrontal and medial temporal cortices). Nevertheless, all within-network correlations were greater than the average across-network correlations for the 17-network parcellation [*t*(31)>4.72, *P*<.0001 for all 17 comparisons]. Given the prominent leveling-off points in the scree plots of Figure 3A and B (right panels) at 3 and 7 - and to lesser extent 13 and 18, the existence of stable parcellations at N=2, 7, and 17 raise the possibility that the scree plots reflect markers of hierarchical network organization.

**Figure 4.**
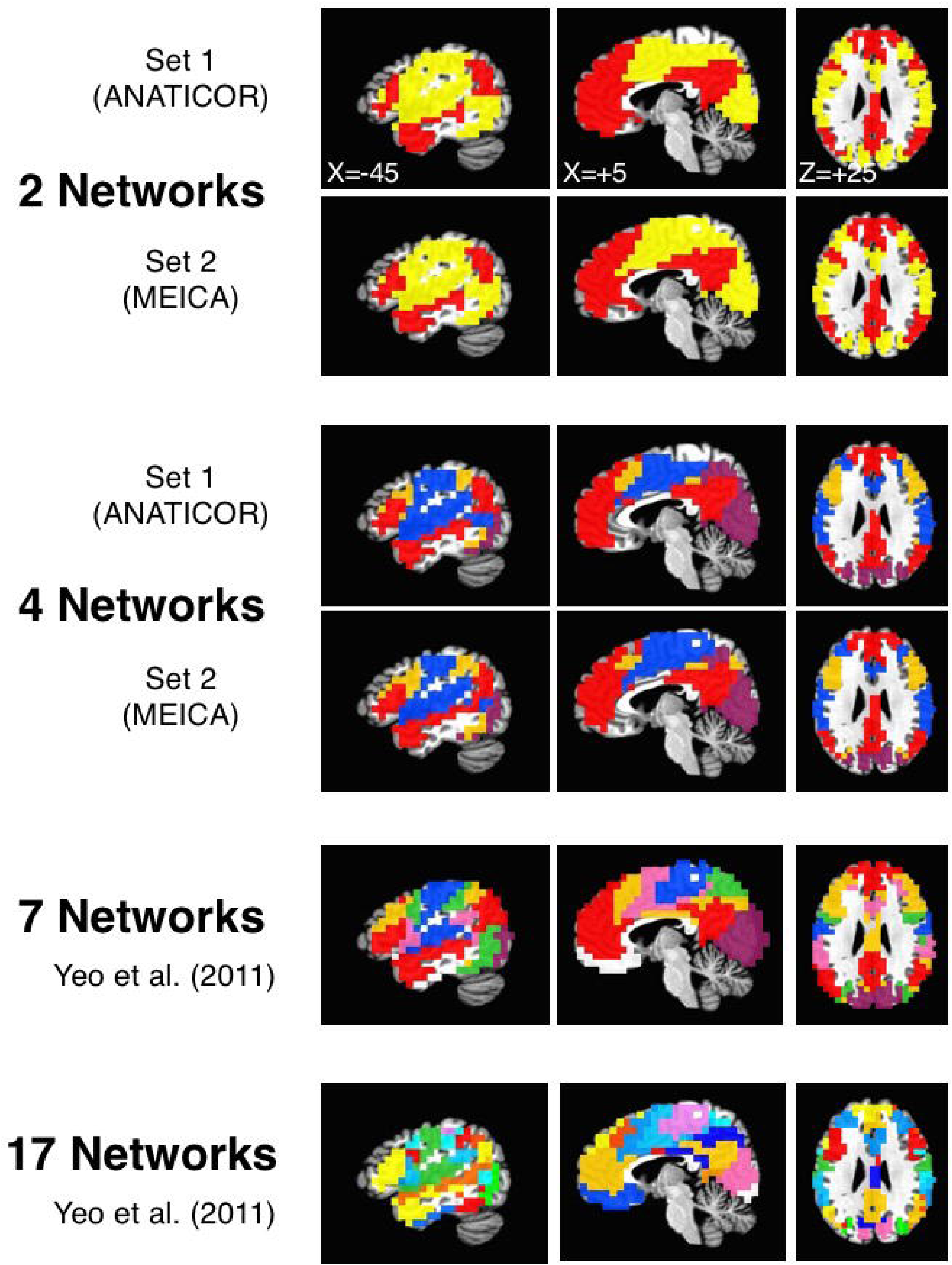
Network parcellations from coarse to fine. The voxelwise correlation matrices calculated on the de-noised data for Sets 1 and 2 (using ANATICOR and MEICA, respectively) were parcellated into networks over a range of thresholds using both Louvain Modularity and InfoMap algorithms. At the lowest thresholds (top two rows), 2 large networks were stable across sets, corresponding to previously described “task positive” (yellow) and “task negative” (red) brain regions (shown uses top 40% of correlation values and Louvain Modularity with gamma=0.75). At higher thresholds, 4 networks were stable in both sets (rows 3 and 4; shown uses top 5% of correlation values and Louvain Modularity with gamma=0.75). Red corresponds to larger DMN, purple corresponds to the Visual network, blue corresponds to the larger Somatomotor network, and orange corresponds to the larger Fronto-parietal network. The fifth and sixth rows show the previously published 7- and 17-network parcellations of Yeo et al. (2011) after first transforming from MNI to Talairach space.

We next examined whether these lower-order parcellations into large networks corresponded to visible clusters in two-dimensional renderings of the unthresholded voxelwise correlation matrix using MDS (as in Figure 2B for *Simulation 2*). For these analyses, the parcellations performed on the de-noised data were applied to MDS analyses of the correlation matrix for the volume registered data. In other words, is this network structure prominent in the data prior to data cleaning when the GS is calculated? Figure 5 shows the results of MDS on the group-average voxelwise correlation matrices for Set 1 in the left column and Set 2 in the right column, with the scatterplots in the top row showing individual black dots for each voxel’s column in the matrix. Nearby dots indicate highly similar patterns of correlation down the rows of the matrix, whereas dots far away indicate dissimilar patterns. The 2- and 4-network parcellations across the Sets 1 and 2 were unified into single sets by retaining only those voxels with identical network membership in both sets. Overall, the correlation structure using MDS was highly similar between the two sets, with one cluster of highly dense points in the lower right of the scatterplots and 1-2 clusters of highly dense points in the lower and upper left. Using the same color schemes as for Figure 4 to show network membership, the 2-network parcellation clearly corresponds to the division between the lower right density of points (task-negative voxels shown in red) and the upper and lower left densities (task-positive voxels shown in yellow). As one progresses to 4 networks, the lower right scatterplot density persists as the task-negative or default-mode network (DMN, in red), the two separate densities on the left correspond mainly to the visual (purple) and somatomotor (blue) networks, and the frontoparietal network (orange) is a mix of both task-positive and -negative voxels with slightly more weighting toward the task-positive voxels. Further differentiation occurs for the 7-network parcellation of Yeo et al. (2011), with the DMN dividing into a continuing DMN (red) and a limbic network (shown in black to be visible against a white background), the fronto-parietal network dividing into a continuing fronto-parietal network (orange) and the dorsal attention network (green), and the somatomotor network dividing into a continuing somatomotor network (blue) and the ventral attention network (pink) (the visual network remains a single network, purple). However, these further networks have less correspondence to clear separate point densities in these two-dimensional scatterplots (with the exception of the limbic network, to the upper right), but rather as adjacent and partly overlapping portions of the previous densities. The best visible agreement between the two approaches appears to occur for the 2-4 network parcellations, and this lower-dimensional network structure is clearly apparent in the MDS results of the volume registered data.

**Figure 5.**
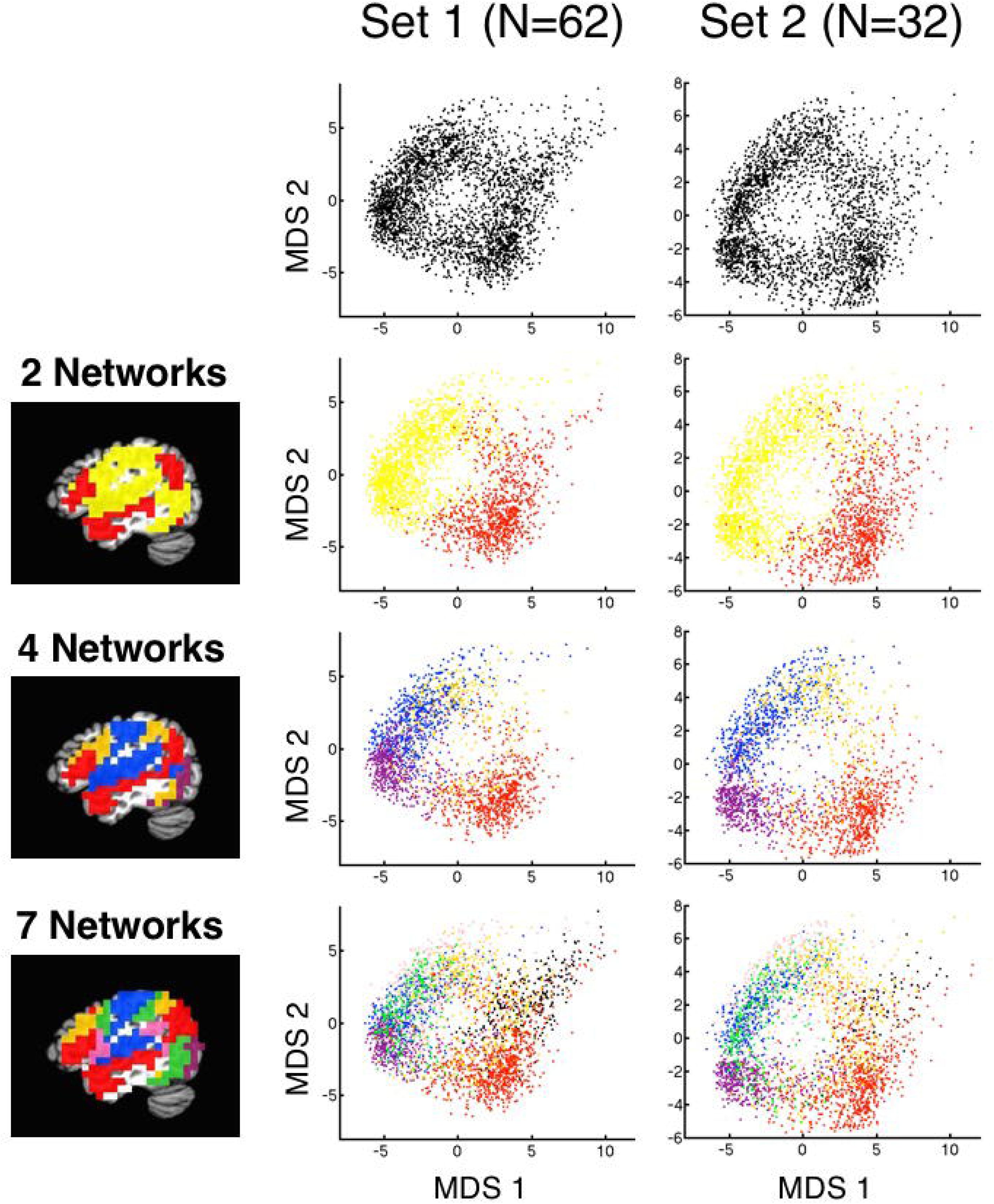
MDS scatterplots shown in relation to network parcellations. The unthresholded group-average voxelwise correlation matrices calculated on the volume registered data were converted to distances and submitted to metric MDS analyses with two dimensions (Set 1 in left column, Set 2 in right column). The top row shows scatterplots for all voxels in the field of view of the Yeo et al. (2011) parcellations, using black dots to represent each voxel’s column in the all-to-all distance matrices. The 2- and 4-network parcellations show results by network using color for voxels that agree in their network classification. The 7-network parcellation of Yeo et al. (2011) is shown in the bottom row, with the limbic network (white in the key) rendered as black dots for visibility on a white background. Viewing the progression from 2 to 7 networks provides a concrete rendition of the hierarchical arrangement of the networks at the voxel level, with high dot densities corresponding best visually to the top levels of the hierarchical tree (2 and 4 networks).

Finally, given the agreement of the network clustering and MDS analyses, we decided to examine the hierarchical structure of the volume registered data using more direct hierarchical clustering analyses for the two datasets (as in Figure 2C for *Simulation 2*). For these analyses, we first extracted network-averaged time series across related voxels for each subject for the 7- and 17-network parcellations of Yeo et al. (2011) using the volume-registered data. We then calculated 7×7 and 17×17 network-level correlation matrices for each subject, averaging across subjects within each set to arrive at the group-level network correlation matrices. The group-average correlation matrices were then converted to Euclidean distances and submitted to hierarchical clustering analysis, with the results for each set viewed using dendrograms (see Methods for details). The 7-network parcellation yielded identical results for the two sets and these agreed well with the results of the cluster analyses across tie density thresholds (Figure 6). The top branch of the clustering trees corresponded to the task-positive/task-negative division. The task-positive branch was then further divided into a larger somatomotor branch and a branch containing the visual network and a larger fronto-parietal network (i.e. the 4-network solution). Finally, these were further divided into the 7-network solution as already described above. The right-most location of the fronto-parietal network within the task-positive branch also reflects its relative similarity to the task-negative network. The hierarchical clustering analysis of the 17-network parcellation did not yield identical solutions across the two sets. However, a best-match analysis did yield consistent results when each of the 17 networks were placed into a “most-similar” branch of the 7-network solution. The synthesized full hierarchical tree spanning the 2- and 17-network levels that replicated across Set 1 and Set 2 is shown in Figure S3. Note the broad similarity of this hierarchical organization with that described earlier by Doucet et al. (2011) when using ICA-based methods, agreeing particularly well in the highest branches of the tree (N=2 through 4 or 5).

**Figure 6.**
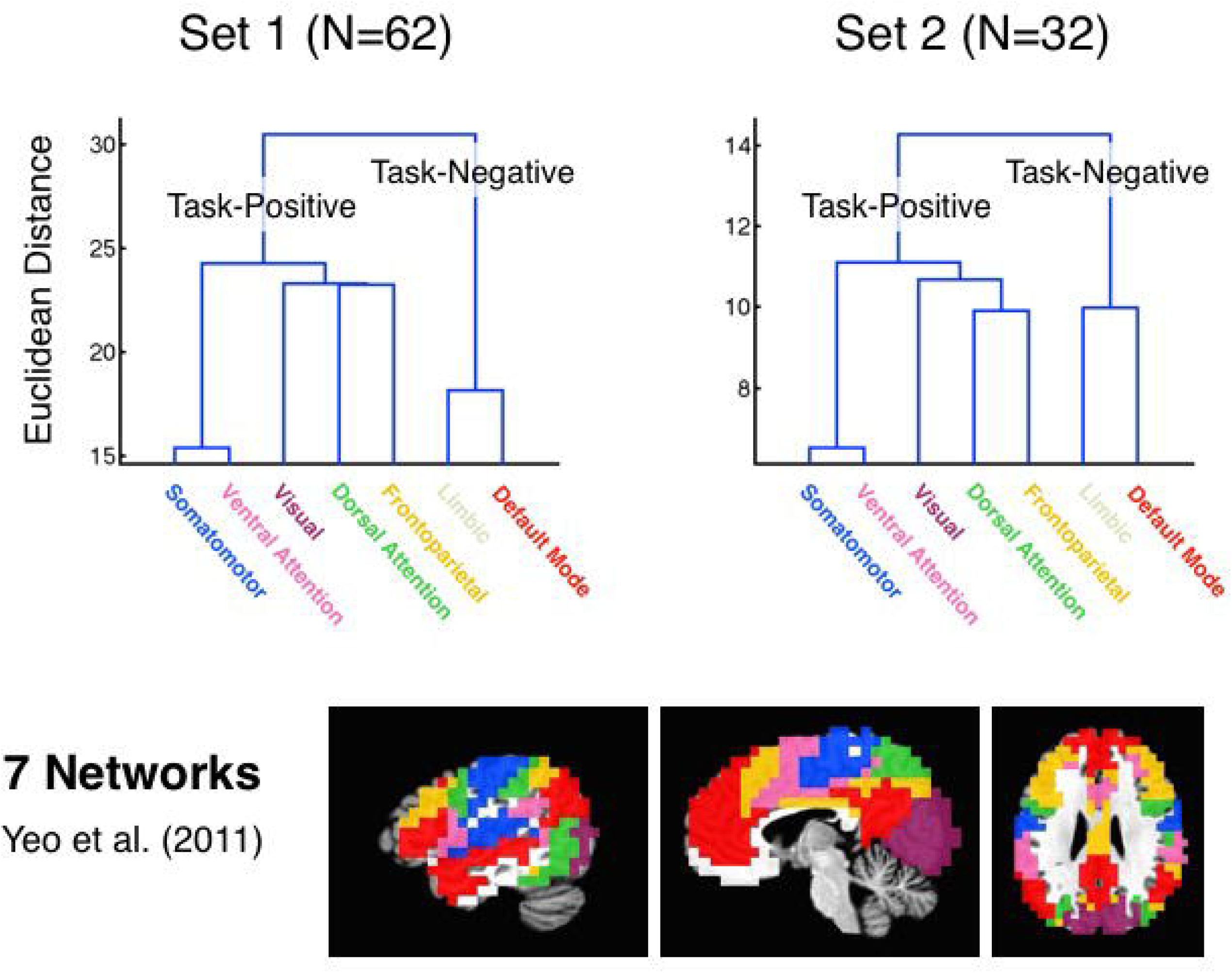
Hierarchical clustering of 7-Network parcellation. Dendrograms based on the network-level correlation matrices of the Yeo et al. (2011) 7-network parcellation and calculated on the volume registered data are shown for Set 1 on the left and Set 2 on the right. Task-positive and task-negative branches divided first, followed by separation of the task-positive branch into broad somatomotor, broad fronto-parietal and visual subbranches, and finally into the 7-network solution. Labels and colors are taken from Yeo et al. (2011).

### Relating the PCA and network clustering analyses

From the previous analyses, it is clear that there are prominent low-dimensional aspects to the voxelwise correlation structure in both the de-noised and the volume-registered data. However, it is still unclear how the information from PCA relates to the different levels of the network hierarchy observed through network clustering. How do different ranges of the PCs relate to specific networks? Is all of the fine-grained information in the 17-network solution contained in PCs 10 to 20? To examine this issue in more depth, we used PCA to reconstruct the voxel-wise time series for each individual subject using different numbers of PCs from 1 to 100. These reconstructed single-subject time series data could then be used to calculate single-subject and group-level voxelwise correlation matrices.

The resulting group-level, voxelwise correlation matrices for the volume-registered data are shown in Figure 7A, with the top row showing results for Set 1 and the bottom row for Set 2. For ease in examining the network relationships, the rows and columns of the matrices have been sorted by the hierarchical network organization shown in Figure S3 (yellow/red brackets next to the axes denote the 2-network parcellation, with the 7-network parcellation labelled in colored rectangles and key shown to the right, and the 17-network parcellation marked in the interior of the matrices with dashed lines). The correlation matrices when using the original full time series of the volume-registered data are shown in the right-most column of Figure 7A for reference. When only including the first two PCs in the time series reconstruction (left-most column of Figure 7A), the main division between task-positive and task-negative is prominent in both datasets, along with some additional network information for the 7- and 17-network parcellations. When using PCs 1-10 (middle column), the patterns present in the correlation matrices are already virtually identical to those when using the full time series. This is quantified in Figure 7B through the use of *R*^2^. When using just the first 2 PCs for reconstruction, the correlation matrices share approximately 60-70% of the variance with the matrix when using the full time series. By the time PCs 1-10 are used, the *R*^2^ is already at approximately 0.9 (90% of the variance). This suggests that the information about even fine-grained networks in the 17-network parcellation are present in PCs 1-10, consistent with a situation of the sort presented in *Simulation 2*.

**Figure 7.**
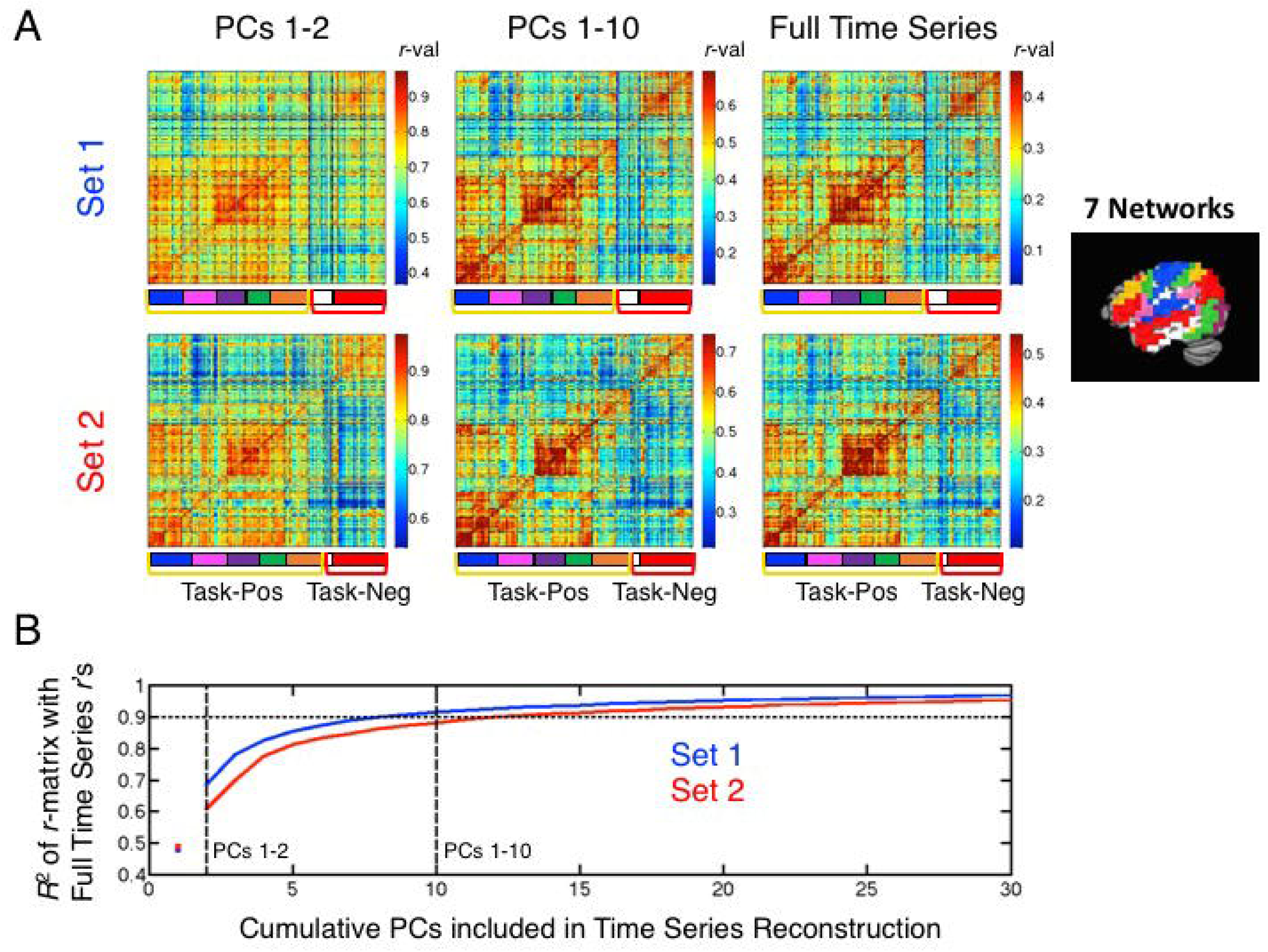
Network information present in the first few PCs. (A) Group-average correlation matrices based on time series reconstructions (volume registered data) and using only a subset of the full PCs are shown for Sets 1 and 2 (top and bottom rows, respectively). Matrix rows/columns have been sorted to reflect the hierarchical arrangement of the full set of network relationships (see Figure S3). Colored brackets (yellow/red) mark the task-positive/task-negative boundaries, with the Yeo et al. (2011) 7-network parcellation boundaries marked with color labels (key shown to the right) and the 17-network parcellation boundaries marked by dashed black lines in the field of the matrices. Reconstructions using only the first two PCs are shown in the left column, using the first 10 PCs in the middle column, and using all PCs (i.e. the full time series) shown in the right column for reference. Correlation values are depicted in the matrices using color, with colorbars shown to the right of each matrix and the max/min color values set as the mean correlation +/− 2 standard deviations to enhance visualization of the network distinctions present in the data. (B) *R^2^* values calculated for the upper triangle of the reconstructed group-average correlation matrices relative to the group-average correlation matrix based on the full time series. The x-axis represents how many cumulative PCs are used in the time series reconstruction (e.g. PCs 1-2 or PCs 1-10), and blue and red curves show the results for Sets 1 and 2, respectively. Values were estimated at PC=1 (shown with a red and blue colored dot) by calculating the *R^2^* for data reconstructed without the 1st PC (i.e. using all PCs but the first) and subtracting from 1. Both curves indicate that approximately 10 PCs are sufficient to explain 90% of the total variance present in the group-average voxelwise correlation matrices calculated using the full time series.

To further examine the fine-grained network information present in the first 10 PCs, we calculated how discriminable the 17 networks were from each other using two measures. We first calculated the number of within- versus across-network correlation comparisons that were significant at *P*<.05, 2-tailed (i.e. is the correlation level within network 1 greater than that between network 1 and network 2) out of a total of 272 possible combinations. We next calculated the mean *t*-value across the related comparisons for a more continuous measure of network discriminability. For both measures, we considered results relative to a baseline of discriminability when using the original full time series. When using the full time series, 92.6% (252/272) of the pairwise within- versus across-network comparisons were significant at *P*<.05 for Set 1, with a mean (SD) *t*-value across comparisons of 6.48 (3.53) with 61 degrees of freedom (for N=62 subjects). For Set 2 when using the full time series, these numbers were 96.3% (262/272) significant at *P*<.05 and a mean (SD) *t*-value of 6.13 (2.93) with 31 degrees of freedom. When including the PCs 1-10 in the time series reconstructions, 249/272 (91.5%) within- versus between-network comparisons were significant at *P*<.05 for Set 1 (98.8% of the level obtained when utilizing the full time series), and 263/272 (96.7%) were significant at *P*<.05 for Set 2 (100% of the level of the full time series). The mean *t*-values were also equal to or slightly greater than the level when using the full time series for both Sets 1 and 2 [mean (SD) for Set 1: 6.48 (3.70); for Set2: 6.39 (2.88); 100% and 104% of level at full time series, respectively]. Indeed, if one asks how many PCs are needed in the reconstruction to achieve 90% of the level of discriminability among the 17 networks for both measures when using the full time series, only PCs 1-7 are needed for Set 1 (238/272, 94.4% of full time series level; mean *t*-value=5.85, 90.4% of full time series level) and PCs 1-6 are needed for Set 2 (253/272, 96.6% of full time series level; mean *t*-value=5.57, 90.9% of full time series level). Taken together, these results show that the vast majority of the information about even the fine-grained networks in the 17-network parcellation are contained in just the first 6-7 PCs.

### Partial correlation analyses of network-specific variance present in the GS

Both the PCA and network clustering results indicate that much of the information about previously reported parcellations - even fine-grained ones - is effectively low dimensional and contained in the first few PCs. This is due to the hierarchical nature of the overall network structure, with a small number of large network branches (N=2 at the top level of the hierarchy) that would not be expected to cancel when averaged. This expectation amounts to a clear prediction: If a small number of mutually orthogonal patterns dominate the average, one should be able to find network-specific variance still present in the GS time series. To examine this prediction, we used partial correlation analyses to check for network-specific variance present in the GS at each level of the network hierarchy from N=2 to N=17. For each subject, network-specific time series were calculated by averaging the time series across voxels within the network. Network-specific time series were derived using the de-noised time series data for each set in order to minimize the amount that noise sources might contribute to the estimates. To estimate a network’s unique contribution to the GS of the volume-registered data, the other network time series at the same level were partialled from that network’s time series and from the GS, calculating the partial correlation coefficient as the Pearson correlation between the two residual time series. This means that any variance shared among the networks due to either neurogenic or artifactual sources were removed from the estimates, leaving only the unique variation shared in the time series due to that specific network. Partial *r*-values were then tested at the group-level against a value of 0 for Sets 1 and 2, with multiple comparisons corrected by false discovery rate (FDR, e.g. Genovese et al., 2002). Box plots conveying information about the full distributions of partial *r*-values across subjects are shown in the top two rows of Figure 8, with the bottom row showing which brain networks replicate in contributing unique variance to the GS across Sets 1 and 2 (green brain masks). For the 2-network parcellation (left column of Figure 8), both task-positive and task-negative network time series contributed unique variance to the GS (Set 1: *P*<3.27×10^−26^ for both; Set 2: *P*<2.65×l0^−15^ for both). For the 4-network parcellation, all four networks contributed unique variance to the GS in Set 1 (*P*<5.41×10^17^ for all) and three out of four contributed significant variance in Set 2 (*P*<5.73<10^10^ for the DMN, fronto-parietal and visual networks), with only the Somatomotor network failing to reach FDR-corrected significance [*t*(31)=1.82, *P*<.079, a non-significant trend]. For the 7-network parcellation, all 7 networks contributed unique variance to the GS in Set 1 (*P*<1.36×l0^−9^ for all), whereas 3 out of 7 were found to contribute unique variance in Set 2 (*P*<1.4×10^−08^ for the DMN, fronto-parietal and visual networks), similar to the results for the 4-network parcellation. Finally, for the 17-network parcellation, 13/17 networks contributed unique variance to the GS in Set 1 (*P*<.0015 for all 13), with one additional network contributing an an uncorrected level [network 16: *t*(61)=2.92, *P*<.005]. In contrast, 3/17 networks (networks 1, 11, and 17) contributed unique variance to the GS in Set 2 (*P*<.0015 for all 3), with networks 3 and 12 contributing at an uncorrected level (*P*<.05 and *P*<.0066, respectively) and networks 7 and 16 showing non-significant trends (P=.054 and *P*=.074, respectively). While only 3/17 networks replicated across Sets 1 and 2, Set 2 had less statistical power to show such effects (N=32 compared to N=62 for Set 1). If one considers effects that showed at least a trend in both sets, 7/17 demonstrated some unique contribution to the GS. In any case, consistent with the prediction of low dimensionality, multiple networks were found to contribute unique variance to the GS at each level of the network hierarchy. This picture did not vary qualitatively as a function of how the data were de-noised.

**Figure 8.**
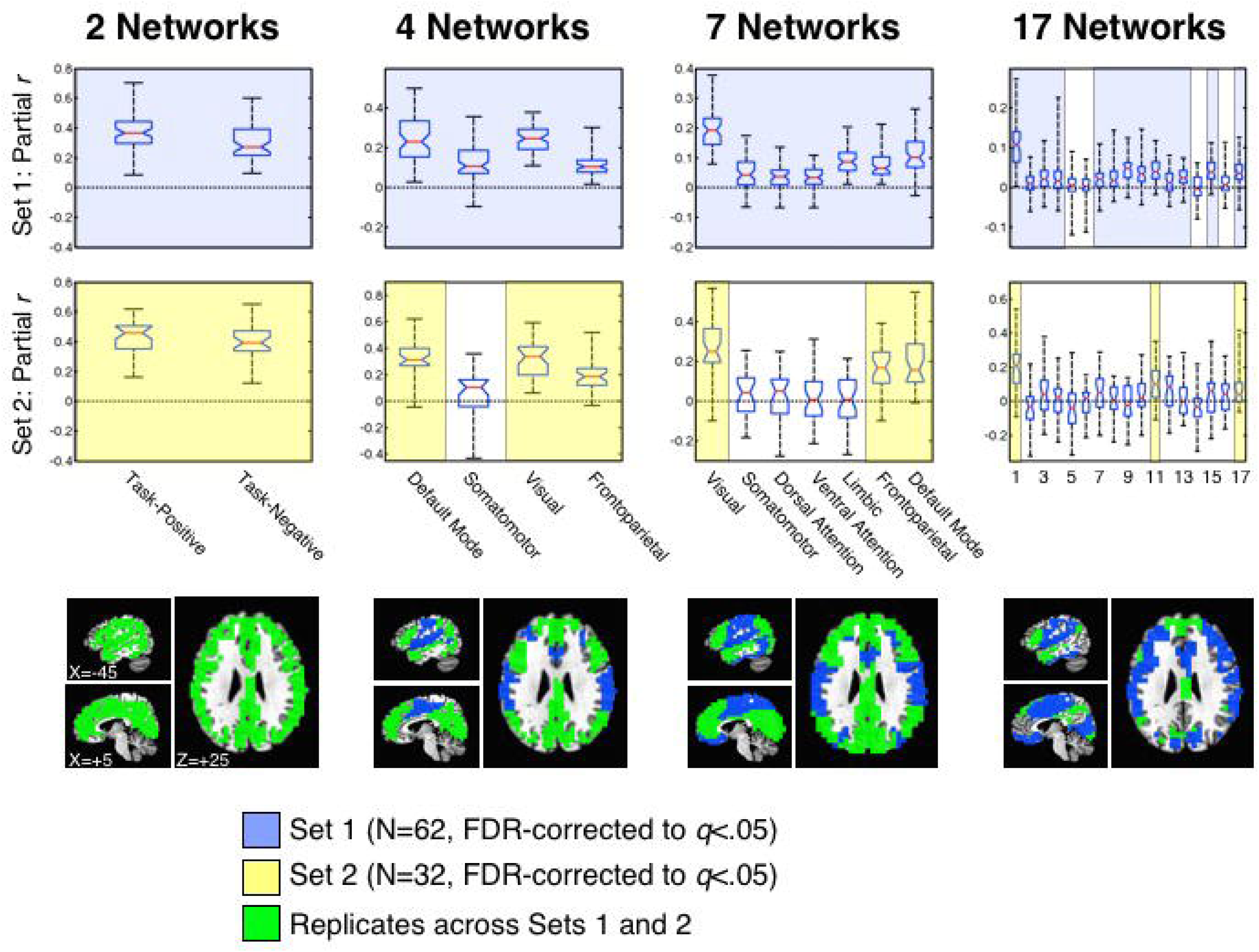
Partial correlation analyses of network-specific information present in the GS. Shown are box plots of the partial *r*-values of each network with the GS, removing shared variance with the other networks at the same level of the network hierarchy. The red horizontal line in each box plot represents the median (50th %-ile), the blue horizontal lines just above and below the median represent the 25th and 75th %-iles, the black horizontal lines represent the minimum and maximum values, and the boundaries of the horizontal notches inside the 25th and 75th %-iles depict the 95% confidence limits of the median. The different network parcellations are shown across the columns (N=2, 4, 7, and 17), the first row shows results for Set 1, and the second row shows results for Set 2. All partial *r*-values were tested against 0 at the group level using one-sample *t*-tests (after applying Fisher’s z-transform to improve normality), correcting for multiple comparisons using FDR. Comparisons surviving FDR correction are highlighted with light blue for Set 1 and light yellow for Set 2. The bottom row renders the locations of the networks that survive correction and replicate across the two sets (shown in green). Network labels/codes for the 7- and 17-network parcellations are taken from Yeo et al. (2011) and match the publicly available parcellations found at https://surfer.nmr.mgh.harvard.edu/fswiki/CorticalParcellation Yeo 2011.

Out of concern that residual global artifacts might somehow contribute variance to the partial-*r* estimates, we conducted a set of follow-up analyses to examine whether the partial correlation approach was removing common variance as expected. It is now well established that even after de-noising, residual head motion and other artifacts can remain in resting-state data (see Power et al., 2015; Satterthwaite et al., 2017, for reviews). We took each subject’s network-averaged time series in the de-noised data and simply regressed it against the GS time series either prior to partialling the other networks or afterwards (other networks were only partialled from the network time series for these analyses rather than from the GS, too, leaving the common variance in the GS). We then examined the beta coefficients from these regressions against the degree of head motion that each subject exhibited during the rest scan as measured by AFNI’s function @ldDiffMag in units of mm/TR (comparable to mean Framewise Displacement, Power et al., 2012). If a subject moved a lot during a scan, one expects those artifacts to be present to an extent in both in the individual network time series in the de-noised data and in the GS of the volume registered data, inflating the beta coefficient. For a subject who moved less, there should be less inflation of the same beta coefficient. After partialling common variance from the networks, the inflation of the beta coefficients by head motion should be effectively removed since that variance is highly similar across the brain. In accord with these expectations, there were strong residual effects of head motion on the network->GS beta coefficients in Set 1 prior to partialling (cleaned by ANATICOR), with correlations ranging from 0.408 to 0.683 [mean=0.587, SD=0.069] across all network time series and network levels examined (all FDR-corrected with *P*<.001). After partialling the other network time series, the network->GS beta coefficients no longer significantly correlated with head motion magnitude [mean=0.068, SD=0.178, none surviving FDR correction]. For Set 2, cleaned by MEICA, no network->GS relationships were significantly correlated with head motion magnitude either before [mean=0.245, SD=0.119, none surviving FDR correction] or after partialling other networks [mean=0.032, SD=0.166, none surviving FDR correction]. These results suggest that MEICA may do a better job removing head motion artifacts than ANATICOR (see Kundu et al., 2013; Power et al., submitted, for further discussion). Nevertheless, for the purposes of the current study, the partialling of other network time series does indeed appear to remove residual global artifacts from the partial correlation estimates quite well.

### Amount of variance in the GS explained by network-specific sources

Having established that network-specific variance is present in the GS, it is still unclear whether this amounts to a large portion of variance. If the network-specific variance only makes up 1-2 percentage points of the total GS variance, then GS removal might still not be expected to have a large impact on the resultant signals after de-noising. To approach this question, we used the partial *r*-values from the analyses in Figure 8 to estimate *R*^2^ of the network-specific portions of variance relative to the total GS time series variance for each subject (see Methods for complete details). Given that partialling network time series from other networks within the same large branch of the hierarchy will remove the shared variance that defines that branch, the best estimate of the total network-specific variance will come from the highest level of the hierarchy, namely the division between task-positive and task-negative voxels for the 2-network parcellation. Indeed, the unique signatures from the more specific subnetworks within each large branch will be present in the average of each of the two branches.

The full distributions across subjects for the network-specific *R*^2^ estimates are shown in box plots in Figure 9A, for Set 1 on the left and Set 2 on the right. Medians are shown with horizontal red lines, the 25th and 75th %-iles are marked by the vertical boundaries of the blue boxes, and 95% confidence limits of the medians are rendered using notches. Minimum and maximum values across the populations are indicated with black horizontal bars connected by dashed lines. The mean (SD) *R*^2^ estimates for the task-positive branch of the network hierarchy are 0.109 (0.066) for Set 1 and 0.107 (0.057) for Set 2. For the task-negative branch, the mean (SD) *R*^2^ estimates are 0.070 (0.055) for Set 1 and 0.091 (0.069) for Set 2. When combined into a total estimate of the variance explained in the GS time series by network-specific sources, the mean (SD) estimates are 0.179 (0.116) for Set 1 and 0.198 (0.099) for Set 2. These estimates indicate that at least 18-20% of the GS time series is due to network-specific sources in both datasets on the average. To the extent that neurogenic sources of variance that are common across networks (e.g. Schölvinck et al., 2010; Wen & Liu, 2016) have been removed from these analyses by the partialling step, we would expect the full neurogenic portion of the GS to be even higher, although it is hard to estimate exactly how much higher given the difficulty in separating this common variance from artifact sources. One should therefore think about these estimates as a floor number rather than a ceiling.

**Figure 9.**
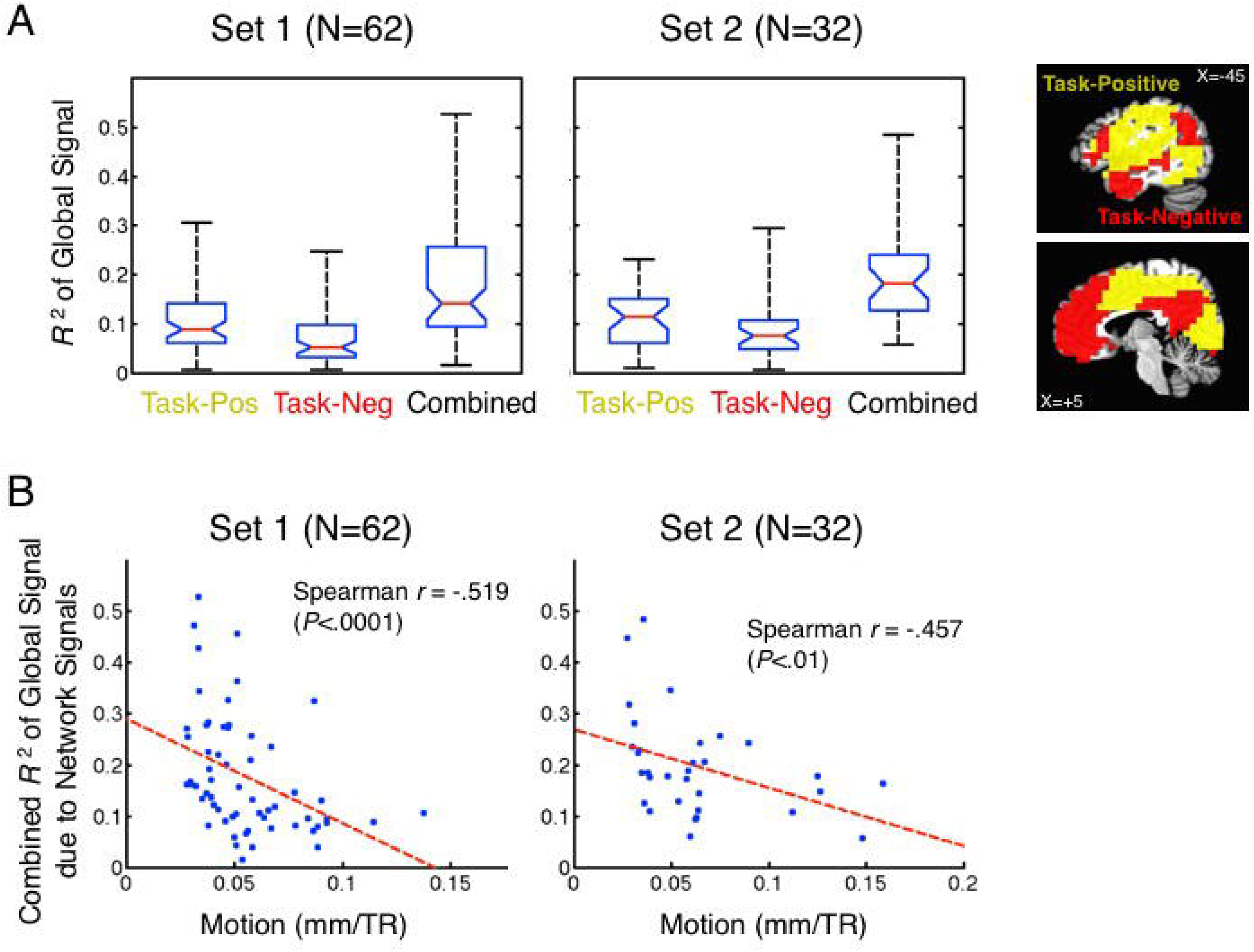
Proportion of variance in the GS due to network-specific sources. (A) Box plots of the *R*^2^ values derived from the partial correlation analyses are shown separately for the N=2 network parcels (Task-Positive, Task-Negative), as well as combined across networks, with Set 1 shown on the left and Set 2 shown on the right. Box plots have the same conventions as described in Figure 8. (B) Dependence on head motion (AFNI’s @1dDiffMag) of the combined *R*^2^ estimates from (A) are shown in scatterplots for Set 1 on the left and Set 2 on the right. In both sets, significant correlations (Spearman) were observed, with lower *R*^2^ values for subjects with higher levels of head motion. Best-fit linear regression lines are shown as dashed red lines.

When examining the individual variability of the single-subject *R*^2^ estimates around the mean in both datasets, it is clear that some values are extremely high (e.g. 0.5 indicating 50% of the variance) whereas some are lower (e.g. 0.1 or below). What might mitigate such a wide range of estimates? One obvious factor to consider is the degree of head motion. For subjects with higher magnitudes of head motion, one can anticipate that a larger fraction of the GS will be due to motion-related signals, resulting in smaller *R*^2^ estimates for the network-specific contributions. Figure 9B shows that this is indeed the case for both Sets 1 and 2, with larger head motion resulting in a smaller *R*^2^ estimate for the network-related signals [Set 1: Spearman *r*(60) = -.519, *P*<.0001; Set 2: Spearman *r*(30) = -.457, *P*<.01]. For subjects with higher head motion, the network-specific estimates shrink from the mean of 20% down to around 10% of the variance in the GS, whereas they increase to 25-30% for subjects with low head motion - with some subjects in both sets having estimates as high as 40-50%. In other words, the subjects with the best data quality can have very large portions of the GS due to network-specific sources. It follows then that using GS regression to de-noise the subjects with the worst data quality comes at the cost of maximally distorting the data from the subjects with the best data quality.

## DISCUSSION

In the current paper, we have examined the dimensionality of resting-state fMRI data in two separate datasets with an eye toward the GS regression debate. We find that while many finegrained brain networks can be distinguished in resting-state data, most of the time series variance related to previously published network parcellations is contained within the first 6-7 PCs. Rather than representing mutually orthogonal time series, these fine-grained brain networks are positioned within a much lower-dimensional network hierarchy (see also Doucet et al., 2011), with the largest branches representing the well-known distinction between task-positive and task-negative -- or alternatively “non-default” and “default” -- brain regions (Buckner et al., 2008; Fox et al., 2005; 2009; Raichle 2015b). The implication of this hierarchical organization, which was illustrated here in simulated as well as real data, is that intercorrelated time series within the large branches do not cancel to 0 when averaged together. Consistent with this, we found that network-specific time series accounted for at least 18-20% of the GS time series variance in both datasets examined. Subjects with the best quality data (low head motion) were found to contribute an even higher proportion of network-specific variance to the GS, reaching as high as 40-50%.

At the core of this issue is the point that networks that are distinct - meaning that time series are more intercorrelated within than across networks - are not necessarily mutually orthogonal. They may simply differ in their level of non-zero (and positive) correlation. When correlation matrices are highly thresholded, the picture that one gets of network number is that the brain is composed of a great many parts with at least some distinct and unique variance. We do not dispute this point at all. Indeed, one could conceptualize the 1000 random time series in *Simulation 1* as describing 1000 mutually orthogonal network time series (for the Zero Clusters case). A full understanding of the brain and its organization would surely take interest in all of this complexity. However, low-order inter-relationships among networks that exist below the thresholding can radically affect the global average of these networks. Adding just two shared time series in *Simulation 1* took the structure from having extremely high dimensionality (of approximately 200 orthogonal dimensions) to having a dimensionality of 2 in the global average. Similarly, in *Simulation 2*, one could describe the number of networks as 25 if one thresholds the all-to-all correlation matrix above an *r*-value of approximately 0.5. However, the network structure below such a threshold has a large impact on the average. Our interpretation of the dimensionality of the actual resting-state fMRI data is that the lower-order correlation structure, described broadly before (e.g. Doucet et al., 2011; Fox et al., 2005), has a similar impact on the actual GS. We believe that the understandable focus on the fine-grained network structure present in the brain - and thinking of these numerous networks as somehow “independent” - has helped lead to expectations of high dimensionality. The findings in the current study highlight the role that the hierarchical nature of such brain networks has in reducing the dimensionality to a much lower level when averaging the signals over the whole brain to a GS.

Some previous studies using PCA to study the GS have also led to confusion about the dimensionality of signals contributing to the GS. Carbonell et al. (2011) previously made the case using PCA to study resting-state fMRI data that the GS was a linearly additive signal with no network sources based on the level of correlation between the first PC and GS, which is ˜ 0.97-0.99. Here we have shown that the conclusions based on this criterion are incorrect, both in the simulated data in which the same phenomenon occurs when the majority of variance in the GS is due to network sources (*Simulation 2*) and in the real fMRI data for which we have established that ˜ 20% of the GS variance on average is due to network-specific sources. Further explorations with *Simulation 1* have shown that such a high correlation between the 1st PC and the GS does seem to require the addition of some level of actual common signal (as was present in *Simulation 2*). However, the added common signal can be well short of a majority of the variance in the GS (e.g. ˜ 10-25% of the variance, with 75-90% due to network structure; see *Supplementary Materials* and Figure S4).

The current approach to estimating network-specific contributions to the GS necessarily removes shared variance among the networks through partialling. Some of this shared variance is expected to be due to neurogenic sources based on previous studies using electrophysiological measures to establish that a portion of the GS reflects real neural activity (e.g. Schölvinck et al., 2010; Wen & Liu, 2016). For example, Schölvinck et al. (2010) used simultaneously acquired microelectrode recordings and BOLD data in monkeys to show that LFP power in high (40-80 Hz) and low (2-15 Hz) frequency bands at several different locations (extrastriate, parietal and frontal cortex) was correlated with BOLD fluctuations broadly over the entire cerebral cortex. These values peaked at correlation levels between 0.2 and 0.3, suggesting approximately 4-9% of the variance accounted for in the GS. Some have suggested that this portion of the GS might correspond to factors such as overall arousal level (e.g. Chang et al., 2016; Pisauro et al., 2016; see Liu et al., 2017, for review). Truly global variance is necessarily distinct from the network-specific portions investigated here, which means that the neurogenic portion of GS variance is likely larger than 20% if this factor is included. How much larger for our current scans is unclear because we do not at present have a clean way to separate neurogenic variance shared across networks from head motion, respiration and other artifactual signals (we do not have veridical measures of these artifact signals either). One way, though, to develop an upper rather than a lower bound on the neurogenic variance is to use the de-noised network-level time series to model total variance in the GS without partialling. These measures will be expected to contain residual nuisance variance, which is why this measure would be an upper bound on the neurogenic variance. After performing these regressions for Set 1 and Set 2, we find that the total variance shared between the network-level data and the GS is 40.3% for Set 1 and 58.1% for Set 2 (see *Supplementary Materials* and Figure S5). This means that the neurogenic portion of variance in the GS is unlikely to be larger than 40-60% for an average subject in our study. Note also that while Set 1 had clear residual motion artifacts (see Results section on *Partial correlation analyses of network-specific variance present in the GS*) and we would expect the 40.3% number to be inflated by this, Set 2 failed to show significant head motion dependence and had the higher proportion of GS variance explained by the MEICA-cleaned network signals (58.1%). However, even if one ignores these numbers entirely and sticks with the lower-bound estimate of 20%, this number can be quite a lot higher for those subjects with lower levels of head motion (as high as 40-50%). It is likely then that the quality control criteria for a given study (our lab commonly requires that mean Framewise Displacement is less than 0.2 mm/TR in order to include the data) will influence the neurogenic portion of the GS, with more distortion due to GS removal for the studies with the best quality data.

### Implications for the GS regression debate

We agree with Power, Petersen, and colleagues that removing the GS will be expected to attenuate motion- and breathing-related variance in the residual resting-state fMRI data, since there is undoubtedly a large portion of variance in the GS that is due to these factors (e.g. Power et al., 2017b; see Liu et al., 2017, for review). Based on the findings in the current study, though, we also fully expect that the presence of prominent network-specific signals in the GS will lead to the additional removal of important signals of neurogenic origin and may also inappropriately alter inter-network relationships. This *may* qualitatively alter group and condition comparisons, as well as correlations with external behavioral variables in the manner described by previous studies (Saad et al., 2012; 2013; Gotts et al., 2013b; Hahamy et al., 2014; Yang et al., 2014). However, as with questions about the dimensionality of the data contributing to the GS average, we’d expect the biggest alterations for phenomena that are more “low dimensional”. In other words, if group or condition differences (or neural correlates of a behavioral measure) are spatially quite extensive over the brain, there is more chance that they will be altered by GS regression. For phenomena that are much more focal in nature or only involve a small number of discrete regions, the alterations to the covariance matrices due to GS regression may have reduced consequences, potentially having a trivial impact on the results of a study. Therefore, an important residual question is: Are the neural correlates of condition or behavioral effects low or high dimensional? For several prominent clinical conditions (e.g. autism, schizophrenia), results from group comparisons with careful head-motion matching of groups and other artifact checking indicate that the neural correlates can indeed be spatially extensive. For autism, we have recently published an updated analysis (Ramot et al., 2017, Supplementary Materials) along the lines of Gotts et al. (2012) using approximately twice as much data (56 ASD, 62 Typically Developing, TD, controls) and matching Age and average Framewise Displacement between the groups. These nuisance variables were further covaried using an Analysis of Covariance (ANCOVA) approach rather than ANOVA or simple *t*-tests. While the most prominent ASD/TD differences involved the STS and somatosensory cortex, the results at an FDR-corrected level (*P*<.05, *q*<.03) involved more than half of the brain in some regard (24396/42988 voxels, or 56.8% of the gray matter). These results are in excellent agreement with the large voxelwise analysis of ASD/TD comparisons using the ABIDE database by Cheng et al. (2015) which also matched and covaried average Framewise Displacement (418 ASD, 509 TD subjects; see Cheng et al., 2015, Supplementary Materials for analyses with and without GS removal). Results this spatially extensive are expected to be accompanied by differing fits to the GS (as shown previously in Gotts et al., 2012, Supplementary Materials), as well as by attenuation and/or alteration of group differences and behavioral correlations due to GS regression (e.g. Gotts et al., 2013b; see also Cheng et al., 2015, Supplementary Materials). The critical follow-up question, though, is whether such alterations due to GS regression reflect distortion based on wide-spread differences in neurogenic signals or reflect differences in residual global artifacts that GS regression is intended to remove. This topic is taken up in more detail in the next section.

### Alternative approaches to accounting for head motion and other artifacts

If one doesn’t remove the GS, what options exist for preventing motion or other artifacts from affecting the results of a study? As already mentioned above, one of the main arguments for removing the GS from the data has been that it is necessary in order to fully remove residual global artifacts such as head motion and respiration from the data of individual subjects (e.g. Power et al., 2014; Satterthwaite et al., 2013; Power et al., submitted). Ours and other groups (e.g. Berman et al., 2016; Gotts et al., 2013b; Saad et al., 2013; Yan et al., 2013a; 2013b) have taken a different approach to addressing these artifacts. The GS is not removed, but groups or conditions are matched on prominent artifact measures such as average Framewise Displacement. This has the effect that little or no between-group variance is explainable by the nuisance measure. Additionally, within-group and between-group variance is modeled in the group/condition comparisons by the slopes of the nuisance measure relative to the functional connectivity values across subjects (using ANCOVA or Linear Mixed Effects models). To the extent that the group effects are uncorrelated with the nuisance measure, then this approach typically leads to stronger rather than weaker group effects, since within-group variance is reduced by the covariate modeling while between-group variance is not due to the matching (for further discussion, see Gotts et al., 2013b; Saad et al., 2013).

This approach will be effective at preventing a global artifact from contaminating group comparisons or correlations with behavior if an artifact is adequately measured by the nuisance measure. However, what if an artifact is poorly measured by the nuisance measure? There are several “omnibus” measures of variance/covariance that one can examine to check for unknown additional global artifacts between groups or that covary with a behavioral measure. GCOR, or global correlation (Saad et al., 2013; Gotts et al, 2013), is the grand mean of the voxelwise correlation matrix. If two groups differ in global artifacts of any kind that impact the average correlation levels, one would expect differences in this measure. However, this measure also has the disadvantage that spatially diffuse but neurogenic differences in correlation levels will also be removed to an extent. A simple alternative to GCOR is checking the average variances of the voxelwise signals. Any artifact source that adds unique variance to the voxelwise time series will inflate the local time series standard deviations (or variances). One can match and covary this measure, averaged over the brain or applied to particular regions of interest. For illustration, this is shown in the *Supplementary Materials* for the 56 ASD and 62 TD subjects previously published in Ramot et al. (2017). Head-motion and more omnibus measures such as GCOR or average local signal amplitudes have now been matched and/or covaried in multiple studies with little or no attenuation of the magnitude of the effects of interest (e.g. Meoded et al., 2015; Berman et al., 2016; Song et al., 2015; Steel et al., 2016; Stoddard et al., 2016; Zachariou et al., 2017). Indeed, in our earlier study on autism (Gotts et al., 2012), we extensively examined head-motion-matched and covaried results for both group differences and behavior with no alteration in the results. Regions differing in functional connectivity also showed no differences in local signal amplitudes that would be reflective of additional, unmodeled sources of residual global artifacts (see Gotts et al., 2012, Supplementary Table 1). While we strongly endorse the current attempts of many groups to develop improved single-subject artifact removal methods, of which MEICA is one (Kundu et al., 2012; 2013), the current state of affairs does not prevent existing studies from drawing solid conclusions from functional connectivity measures without the removal of the GS.

## METHODS

### Subjects

The data in *Set 1* were comprised of 62 right-handed males (mean age = 21.2 years, SD = 5.1 years) with no history of psychiatric or neurological disorders. Informed assent and consent were obtained from all participants and/or their parent/guardian (participants younger than 18), and the experiment was approved by the NIMH Institutional Review Board (protocol 10-M-0027, clinical trials number NCT01031407). Subsets of these data have been used in several previous studies from our lab, serving as TD controls for our studies of autism (Gotts et al., 2012; 2013b; Plitt et al., 2015; Ramot et al., 2017), lateralization of function (Gotts et al., 2013a), and in the development of preprocessing procedures (Jo et al., 2010; Power et al., 2017b). The data in *Set 2* were comprised of 32 right-handed subjects (20 females, 12 males; mean age 24.8 years, SD = 3.9 years) with no history of psychiatric or neurological disorders. Informed consent was obtained from all participants, and the experiment was approved by the NIMH Institutional Review Board (protocol 93-M-0170, clinical trials number NCT00001360).

### MRI acquisition

The data for Set 1 were acquired on a GE 3.0-Tesla whole-body MRI scanner at the National Institutes of Health Clinical Center NMR Research Facility, using standard imaging procedures. For each subject, a high-resolution T1-weighted anatomical image (magnetization-prepared rapid acquisition with gradient echo, or MPRAGE) was obtained (124 axial slices, 1.2-mm slice thickness, field of view = 24 cm, 224 × 224 acquisition matrix). Spontaneous, slowly fluctuating brain activity was measured during fMRI, using a gradient-echo echo-planar imaging (EPI) series with whole-brain coverage while subjects maintained fixation on a central cross and were instructed to lie still and rest quietly (repetition time, TR = 3500 ms, echo time, TE = 27 ms, flip angle = 90°, 42 axial contiguous interleaved slices per volume, 3.0-mm slice thickness, field of view, FOV = 22 cm, 128 × 128 acquisition matrix, single-voxel volume = 1.7 × 1.7 × 3.0 mm^3^). Each resting scan lasted 8 min 10 s for a total of 140 consecutive whole-brain volumes. All EPI data were evaluated for sharp head motion artifacts, with included scans required to be less than or equal to 0.2 mm/TR using AFNI’s @1dDiffMag function (comparable to mean Framewise Displacement, Power et al., 2012). Independent measures of cardiac and respiration cycles were recorded during the resting scans for later removal (all 62 subjects). A GE 8-channel receive-only head coil was used for all Set 1 scans with an acceleration factor of 2 (ASSET), reducing gradient coil heating during the session.

The data for Set 2 were acquired on a GE MR750 3.0-Tesla scanner with a 32-channel receive-only head coil, also at the NIH Clinical Center NMR Research Facility. As with Set 1, all scans were conducted with an acceleration factor of 2 (ASSET). Each subject received a highresolution Tl-weighted anatomical image (MPRAGE) (172 axial slices, 1.0-mm slice thickness, 1.0 mm^3^ isotropic voxels). Eyes-open rest scans were acquired using a BOLD-contrast sensitive multi-echo echo-planar sequence (TEs = 12.5, 27.7, and 42.9 ms; TR = 2200 ms, flip angle = 75°, 64 × 64 matrix, in-plane resolution = 3.2 × 3.2 mm). Whole-brain EPI volumes of 33 interleaved, 3.5-mm-thick oblique slices (manually aligned to the AC-PC axis) were obtained every 2.2 s. All EPI data were evaluated for sharp head motion artifacts, with included scans required to be less than or equal to 0.2 mm/TR using AFNI’s @1dDiffMag. Independent measures of cardiac and respiration cycles were recorded during the resting scans for later removal/examination, although recording errors (dropped samples) prevented the use of these data in 7/32 subjects.

### MRI preprocessing

#### Single-echo processing

All Set 1 EPI scans were single-echo (TE=27 ms), whereas only the middle echo (TE=27.7 ms) was used to calculate the volume-registered data for Set 2 scans, since this TE was comparable to the appropriate echo time to optimize T2* contrast. First, the initial 3-4 TRs were removed to allow for T1 equilibration (4-5 times the T1 of gray matter or ˜ 8 seconds: 3 TRs for Set 1 and 4 TRs for Set 2). Preprocessing utilized the AFNI software package (Cox, 1996), applying 3dDespike to bound outlying time points per voxel within 4 standard deviations of the time series mean, 3dTshift to adjust for slice acquisition time within each volume (to *t*=0), 3dvolreg to align each volume of the resting-state scan series to the first retained volume. Scans were then spatially blurred by a 6 mm Gaussian kernel (full width at half maximum), divided by the voxelwise time series mean to yield units of percentage signal change, and spatially transformed to standard anatomical space (Talairach-Tournoux) with a downsampled spatial resolution (6mm isotropic voxels) to minimize computational time for later voxelwise PCA and MDS analyses. Finally, low frequency trends were removed using a 4th-order baseline polynomial function (3dDetrend). The GS was calculated for each subject from these re-scaled and detrended volume-registered data as the mean time series within a whole-brain mask.

For data in Set 1, de-noising used the ANATICOR preprocessing approach (Jo et al., 2010; see also Gotts et al., 2012) just prior to Talairach transformation in the above steps. White matter and large ventricle masks were created from the aligned MPRAGE scan using Freesurfer (e.g. Fischl et al., 2002). These masks were resampled to EPI resolution and then eroded by 1 voxel to prevent partial volume effects with gray matter voxels. Nuisance regressors for each voxel included included: 6 head-position parameter time series (3 translation, 3 rotation), 1 average eroded ventricle time series calculated from the blurred and re-scaled volume-registered data, 1 “localized” eroded white matter time series (averaging the time series of all white matter voxels within a 15mm-radius sphere), 8 Retroicor time series (4 cardiac, 4 respiration) calculated from the cardiac and respiratory measures taken during the scan (Glover et al., 2000), and 5 Respiration Volume per Time (RVT) time series to minimize end-tidal C02 effects from deep breaths (Birn et al., 2008). Prior to regression, all time series were detrended by a 4th-order polynomial function to remove slow scanner drift and drift in head position. After regression, cleaned time series were transformed to Talairach space and downsampled to 6mm isotropic voxels, as with the volume-registered data.

#### Multi-echo processing

Multi-echo ICA (MEICA) de-noising on Set 2 data was carried out using the meica.py script that is available with AFNI. The initial steps of meica.py are similar to the steps for the single-echo preprocessing, but applied to all 3 echos of data: removing 1st 4 TRs, despiking the voxel-wise time series (3dDespike), adjusting for slice-time acquisition (3dTshift), rigid-body registration of all volumes to the first retained TR (3dvolreg). All echos of data were then masked (whole-brain) and submitted to tedana.py with default settings, which uses the signal decay properties across echos to separate time series into BOLD-like and non-BOLD-like components derived from Independent Component Analysis (ICA) using automated criteria (see Kundu et al., 2012; 2013, for full details). After MEICA de-noising, BOLD-like data were transformed to Talairach space and downsampled to 6mm isotropic voxels, as with the volume-registered data.

### Principal Component Analysis (PCA)

We applied PCA to the voxelwise volume-registered timeseries data of individual subjects, using Matlab’s *princomp* function. This function returned the coefficient matrix, the component scores (the projection of the data into component space), as well as the eigenvalues of each component (the variances of the component scores). The eigenvalues could then be used to characterize how much variance out of the total each component accounted for in the voxelwise time series data, dividing each eigenvalue by the sum of all eigenvalues. PCA was also applied to the group-average voxelwise covariance/correlation matrix of the de-noised data using the Matlab function *pcacov*, returning the eigenvalues for each component. Time series reconstructions for individual subjects including only the first *X* PC’s was achieved by reversing the projection into component space using the inverse of the coefficient matrix:

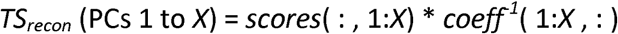

Results were also checked against the Matlab function *pcares*. Following reconstruction, voxelwise correlation matrices could then be calculated for each subject using the new time series, followed by group-average correlation matrices. Comparisons between parcellated networks using reconstructed time series were done by calculating the average within-network level of correlation for each individual subject, as well as the average across-network correlations for each network combination. Statistical comparisons were then calculated across subjects using *t*-tests after first applying the Fisher’s *z*-transform to the correlations to yield more normally distributed values.

### Network parcellations

Group-averaged, voxelwise correlation matrices calculated on the de-noised data for Sets 1 and 2 (using ANATICOR and MEICA, respectively) were thresholded (binary undirected) over a wide range of thresholds and submitted to cluster analyses using the Louvain Modularity (Blondel et al., 2008; Rubinov & Sporns, 2010) and InfoMap (Rosvall & Bergstrom, 2008, 2011) algorithms. Included voxels were the intersection of the scanning field of view for each set and the field of view of the “liberal” gray matter parcellations of Yeo et al. (2011; https://surfer.nmr.mgh.harvard.edu/fswiki/CorticalParcellation Yeo 2011) after first transforming these parcellations to Talairach space to insure the most comparable cohort of voxels. Correlation values were ranked and thresholded in terms of tie density (top *X* percentage) over the whole matrix, using values: top 90%, 80%, 70%, 60%, 50%, 40%, 30%, 20%, 15%, 10%, 9%, 8%, 7%, 6%, 5%, 4%, 3%, 2%, 1%, 0.5%. Louvain Modularity was applied to all of these thresholds for both sets, and InfoMap was applied to all values of top 20% or higher (excluding lower thresholds due to lengthy run times). For Louvain Modularity, a range of the ‘gamma’ parameter (the resolution of the parcellations) was explored, with 1.0 (classic modularity), less than 1.0 (courser: 0.75), and greater than 1.0 (finer: 1.25,1.5, 2.0, 2.5). Each threshold was tested for 100 iterations for Louvain Modularity due to the stochastic aspects of the clustering procedure. From these 100 iterations, a summary matrix was constructed containing the likelihood that each voxel was clustered together with every other voxel (values ranging from 0 to 1). This likelihood matrix was then thresholded at 0.5 and clustered one final time. Cluster solutions using InfoMap were taken as the optimal solution over 1000 iterations. Small and singleton clusters (i.e. a single voxel) derived from either algorithm were excluded by requiring that retained clusters be at least 1% in size of the total number of voxels being clustered (2895 total 6mm^3^ voxels for Set 1; 2700 voxels for Set 2; these differ due to slightly different scanning fields of view in the two sets). Solutions were counted as stable if both sets gave matching cluster solutions at the same or adjacent thresholds when applying the same algorithm.

### Agreement of parcellations and MDS

Multidimensional Scaling analyses (MDS) were applied to the unthresholded group-average voxelwise correlation matrices of Sets 1 and 2 after first converting these matrices to distances using the Matlab function *pdist*. All MDS analyses compressed the full similarity space to two dimensions for ease of viewing, employed the metric version of MDS (Matlab’s *mdscale*), and minimized the squared stress goodness-of-fit criterion. Group-average correlation matrices submitted to MDS on the actual resting-state fMRI data for Sets 1 and 2 were calculated on the volume-registered data using a field of view that overlapped the Yeo et al. (2011) parcellations of gray matter voxels. For the 2-network and 4-network parcellations, only voxels that agreed in network membership across the two sets were included for viewing in MDS scatterplots. Network membership in these scatterplots was conveyed by distinct colors for each network.

### Hierarchical clustering analyses

We examined the hierarchical structure of the volume registered data using hierarchical clustering analyses for Sets 1 and 2. We first extracted network-averaged time series for each network in the 7- and 17-network parcellations of Yeo et al. (2011). We then calculated group-averaged 7×7 and 17×17 network-level correlation matrices and converted these to Euclidean distances (Matlab’s *pdist* function). The distance matrices were then submitted to hierarchical clustering analysis (Matlab’s *linkage* function), with the results for each set viewed using dendrograms. Results were considered stable if the dendrograms had the same branching structure for the two sets.

### Partial correlation analyses and *R*^2^ estimates

We used partial correlation analyses to check for network-specific variance present in the GS at different levels of the network hierarchy. For each subject, network-specific time series were calculated by averaging the time series across voxels within each network for the N=2, N=4, N=7, and N=17 parcellations. This was done on the de-noised time series data for Sets 1 and 2 in order to minimize the amount that noise sources might contribute to the estimates. To estimate a network’s unique contribution to the GS of the volume-registered data, the other network time series at the same level were partialled from that network’s time series and from the GS using least squares multiple regression. The Pearson correlation of these two residual series is equal to the corresponding partial Pearson correlation coefficient (the partial *r*-value). partial *r*-values were then transformed by Fisher’s z to improve normality and were tested at the group-level against a value of 0 for Sets 1 and 2, with multiple comparisons corrected by false discovery rate (FDR, e.g. Genovese et al., 2002) over all individual network tests within a set (e.g. 2+4+7+17=30 total tests for Set 1 or Set 2). Estimates of *R*^2^ were then calculated for each network by squaring the partial *r*-value and multiplying it by the ratio of the residual variance left in the GS after partialling relative to the total variance in the GS prior to partialling (i.e. adjusting for the amount of variance removed by the other networks in terms of the original GS variance). The total *R*^2^ in the original GS due to network-specific sources was then the sum of the individual network *R*^2^ values when using the N=2 parcellation, since using finer-grained parcellations would partial away the shared variance that defines these two large network branches.

The success of the partialling procedure at removing shared global variance due to artifactual sources such as head motion was checked using regression analyses. Each network-specific time series (the predictor variable) was regressed against the original GS (dependent variable) both prior to and after partialling the other network-specific time series from the predictor variable at the same level of the network hierarchy, yielding a beta weight that represented the degree of fit. The choice of beta weights for this purpose is important, since the beta weight does not depend on additional variance present in the dependent variable (the GS) that is uncorrelated with the predictor variable (i.e. the network-specific time series). If shared variance due to factors such as motion is removed from the predictor variable by partialling the other network time series, the remaining presence of motion information in the full GS will then have no residual impact on the beta weight estimate. If Pearson correlation were chosen instead of regression, then the resultant *r*-values would always exhibit motion dependence to the extent that motion variance is present in the GS, since the variance of both variables is used in normalizing the *r*-value. After acquiring the beta estimates, the intersubject variability in the beta weights from these regressions was then examined against the amount of head motion present in the scan (AFNI’s @1dDiffMag, comparable to mean Framewise Displacement over a scan, Power et al., 2012) using Pearson correlation, with multiple comparisons corrected by FDR over all individual network tests within a set (i.e. 2+4+7+17=30 total tests). This approach is similar to quality-control analyses used by other groups (e.g. Satterthwaite et al., 2012; Power et al., 2014) to examine the motion dependence of functional connectivity estimates.

### Simulation 1

In this simulation, 1000 random time series were created by randomly sampling a Gaussian distribution (mean=0, SD=1) using the Matlab function *randn*, each of length 200 time points. In the Two Clusters condition, a simple correlation structure was included by adding a new common random time series to 500 of the original time series (also generated using *randn*) and a different common time series to the remaining 500 original time series. This resulted in two large time series clusters (Cluster 1:1-500, Cluster 2: 501-1000). In the Zero Clusters condition, no shared time series were added. GS time series were created for each condition by averaging over all 1000 time series, and dimensionality and network structure were assessed using PCA and MDS.

### Simulation 2

In this simulation, 1250 random time series were created by randomly sampling a Gaussian distribution (mean=0, SD=1) using the Matlab function *randn*, each of length 200 time points. A nested, 5x5 hierarchical structure was created by adding 5 new common random time series (also using *randn*) to ranges 1-250, 251-500, 501-750, 751-1000, and 1001-1250 and then 25 additional random time series to subranges in groups of 50 (e.g. 1-50, 51-100,101-150,…, 1201-1250). A global common time series was then further added to each of the 1250 time series at half amplitude of the other random series (i.e. scaled by 0.5: Gaussian mean=0, SD=0.5). A GS time series was created by averaging over all 1250 time series, and dimensionality and network structure were assessed using PCA, MDS, and hierarchical clustering (Matlab’s *linkage* function as described above in *Hierarchical clustering analyses*). To evaluate the claims of Carbonell et al. (2011), the simulation was repeated 1000 times, each time calculating the correlation between the first PC and the GS.

## SUPPLEMENTARY MATERIALS

Supplementary materials can be found online at XXX.

## DATA AVAILABILITY

Data are available via the XNAT platform (dataset title ‘XXX’ with ID number XXX). As subjects in Set 1 signed an older version of the consent form, which does not explicitly allow for data sharing, we are currently working on re-consenting all of these subjects with the new version. For now, users will need to request access through the XNAT system. This can be done by creating an XNAT user account and pressing the ‘request access’ link. All of the subjects in Set 2 are currently available for data sharing.

## ACKNOWLEDGEMENTS

The authors would like to thank Drs. Steve Petersen, Jonathan Power, Bob Cox, Ziad Saad, Gang Chen, and Peter Bandettini for helpful discussions. This work was supported by the Intramural Research Program, National Institute of Mental Health (ZIAMH002920).

